# Bifunctionality of a biofilm matrix protein controlled by redox state

**DOI:** 10.1101/138099

**Authors:** Sofia Arnaouteli, Ana Sofia Ferreira, Marieke Schor, Ryan J. Morris, Keith M. Bromley, Jeanyoung K. Jo, Krista L. Cortez, Tetyana Sukhodub, Alan R. Prescott, Lars E.P. Dietrich, Cait E. MacPhee, Nicola R. Stanley-Wall

## Abstract

Biofilms are communities of microbial cells that are encapsulated within a self-produced polymeric matrix. The matrix is critical to the success of biofilms in diverse habitats, but despite this many details of the composition, structure, and function remain enigmatic. Biofilms formed by the Gram-positive bacterium *Bacillus subtilis* depend on the production of the secreted film-forming protein BslA. Here we show that a gradient of electron acceptor availability through the depth of the biofilm gives rise to two distinct functional roles for BslA and that these can be genetically separated through targeted amino acid substitutions. We establish that monomeric BslA is necessary and sufficient to give rise to complex biofilm architecture, while dimerization of BslA is required to render the community hydrophobic. Dimerization of BslA, mediated by disulfide bond formation, depends on two conserved cysteine residues located in the C-terminal region. Our findings demonstrate that bacteria have evolved multiple uses for limited elements in the matrix, allowing for alternative responses in a complex, changing environment.

**Significance:** The biofilm matrix is a critical target in the hunt for novel strategies to destabilise or stabilise biofilms. Knowledge of the processes controlling matrix assembly is therefore an essential prerequisite to exploitation. Here we highlight that the complexity of the biofilm matrix is even higher than anticipated with one matrix component making two independent functional contributions to the community. The influence the protein exerts is dependent on the local environmental properties, providing another dimension to consider during analysis. These findings add to the evidence that bacteria can evolve multifunctional uses for the extracellular matrix components.

## Introduction

Biofilms are assemblies of microbial cells that are attached to a surface or each other (1). Assembly is facilitated by manufacture of an extracellular matrix, which provides both structural and biochemical support to the biofilm community (2). After analysis of many diverse species, the biofilm matrix has been found to mainly comprise exopolysaccharides, extracellular DNA (eDNA) and proteins, many of which form higher order structures such as fibres or filaments (3). Currently there is increasing knowledge of the nature and form of the individual components in the matrix (3) but how the matrix components are deployed in the biofilm, how they interact with other elements in the matrix, and how the local physicochemical environment impacts the properties of the materials used in the biofilm are under-explored.

*Bacillus subtilis* is a Gram-positive, spore forming bacterium, which is found in great abundance in the soil (4) and forms biofilms with a distinctive complex architecture (5). The biofilms display an unusual property: they are highly hydrophobic and remain as such after contact with water, inorganic or organic solvents, and commercial biocides (6). The hydrophobicity of the biofilm has been attributed to a small secreted protein named BslA (7, 8), which works alongside the fibrous protein TasA (9, 10) and the extracellular polysaccharide produced by the products of the *epsA-O* operon (5) to allow morphogenesis of the biofilm. Partial evidence of the mechanism employed by BslA to confer hydrophobicity to *B. subtilis* biofilm has been revealed (8, 11–13). While transcription of *bslA* occurs uniformly within in the population, secreted BslA is localized to the periphery of the biofilm, where it forms a hydrophobic layer at the air-biofilm surface (8, 12). Consistent with this, biophysical analysis of recombinant BslA *in vitro* demonstrated spontaneous formation of a stable elastic protein film at an air-water interface (12). Structural analysis revealed that BslA is an amphipathic protein consisting of an immunoglobin G-like scaffold appended with a hydrophobic “cap” (12) that can present in two forms: ‘cap in’ and ‘cap out’. This property either shields (cap in) or reveals (cap out) the hydrophobic domain in response to the local environment (11). Thus, when BslA is in the aqueous environment of the matrix, it is likely that the hydrophobic cap is ‘hidden’ towards the interior of the protein. In contrast, when BslA reaches a hydrophobic interface (e.g. air at the surface of the biofilm) it undergoes a limited conformational change to expose the hydrophobic cap. Nonetheless, the mechanism by which the BslA biofilm surface layer assembles *in vivo* is undefined.

*B. subtilis* encodes a monomeric BslA paralogue called YweA (14). Deletion of *yweA* does not impact the overall morphology or hydrophobicity of the biofilm (8); however deletion in combination with removal of *bslA* exacerbates the biofilm defect of the single *bslA* deletion (8, 14).Contrary to the marginal contribution of YweA to biofilm formation, but consistent with the high level of amino acid sequence similarity, *in vitro* recombinant YweA undergoes the partial structural rearrangement at an interface to reveal the hydrophobic cap, and forms an elastic protein film, albeit with limited stability (14). One notable difference between the primary amino acid sequences of BslA and YweA is that the BslA-like variants possess a short C-terminal extension that contains two conserved cysteine residues in a “CxC” configuration. Cysteine residues play an important role in the function of diverse proteins in a wide range of cellular processes (15) and therefore we evaluated if the BslA ‘CxC’ motif played a significant biological role or was functionally redundant. Our analysis indicates that in addition to BslA having two structural forms (x0027;cap in’ / ‘cap out’) (11) it also has two *functional* forms mediated not by the hydrophobic cap region but by the cysteines at the C-terminus: monomeric BslA dictates biofilm structure, whereas surface hydrophobicity requires at least dimeric protein. Thus, we show that the cysteine residues are crucial for full BslA function, and surmise that in the native biofilm, electron acceptor availability influences BslA oligomerization: in the oxygen-rich, surface-exposed region, the predominant form is a dimer, while the anoxic environment in the depths of the biofilm prevents BslA dimerization and allows for nutrient uptake. To our knowledge this is the first example of a biofilm matrix protein with two functions that can be genetically separated and where the activity is controlled by redox state.

## Results

### BslA disulfide bond formation

Recombinant BslA_42-181_ forms a mixture of predominantly monomeric and dimeric protein *in vitro*, with a small amount of tetramer observed (12) (Fig S1). The requirement of the cysteine residues (C178 and C180) for oligomerisation of BslA was assessed *in vitro* using purified recombinant protein where the cysteine residue(s) were either replaced with alanine or where the C-terminal 10 amino acids were removed (Fig. S1). Replacement of either C178 or C180 with alanine abolished tetrameric BslA (Fig. S1A and C) but dimers, which could be disrupted by addition of β-mercaptoethanol, still formed (Fig. S1). In contrast, when both cysteines were replaced with alanine, or the C-terminal 10 amino acids were deleted, the protein was restricted to a monomeric form (Fig. S1). Secondary structure analysis of the proteins by circular dichroism (CD) spectroscopy indicated that each of the BslA variants had the same overall fold as the wild-type protein, indicating that the inability to form dimers or tetramers was not due to a disruption of the protein structure (Fig. S2A).

Next, we tested to see if BslA formed dimers and/or tetramers *in vivo*, to explore the possibility that the conserved “CxC” motif at the C-terminus of BslA plays an *in vivo* role in BslA oligomerisation. Proteins were extracted from the wild-type biofilm either in the presence of Cu(II)-(o-phenanthroline)_3_, which cross-links disulfide bonds (16), or 10 mM DTT, a reducing agent. Western blotting revealed immuno-reactive bands consistent with monomeric (~14 kDa), dimeric (~28 kDa) and tetrameric (~ 56 kDa) forms of BslA in the absence of reducing agent (Fig 1A), while in the presence of 10 mM DTT only monomeric BslA was detected (Fig. 1B). The *bslA* deletion strain (NRS2097)was used as a control for antibody specificity, and did not reveal any interacting bands. We generated a series of strains to express the variant *bslA* coding regions in the *B. subtilis bslA* deletion strain (Table S1). Analysis of proteins extracted from the mature biofilms by immunoblot showed the same BslA monomer, dimer and tetramer pattern *in vivo* as that observed *in vitro* for the recombinant proteins (compare Fig. S1 with 1A). Strains generating C178A (NRS5177) and C180A (NRS5178) variant BslA formed dimers but not tetramers, while the double C178A, C180A mutant (NRS5179) and the BsIAai_72-_i_8_i (NRS2957) truncation variants were restricted to the monomeric state (Fig. 1A). In each case only monomeric BslA was detected by Western blot analysis when 10 mM DTT was added during protein extraction (Fig. 1B). These data demonstrate that BslA forms dimers and tetramers *in vivo* in a C178 and C180 dependent manner.

**Figure 1:**
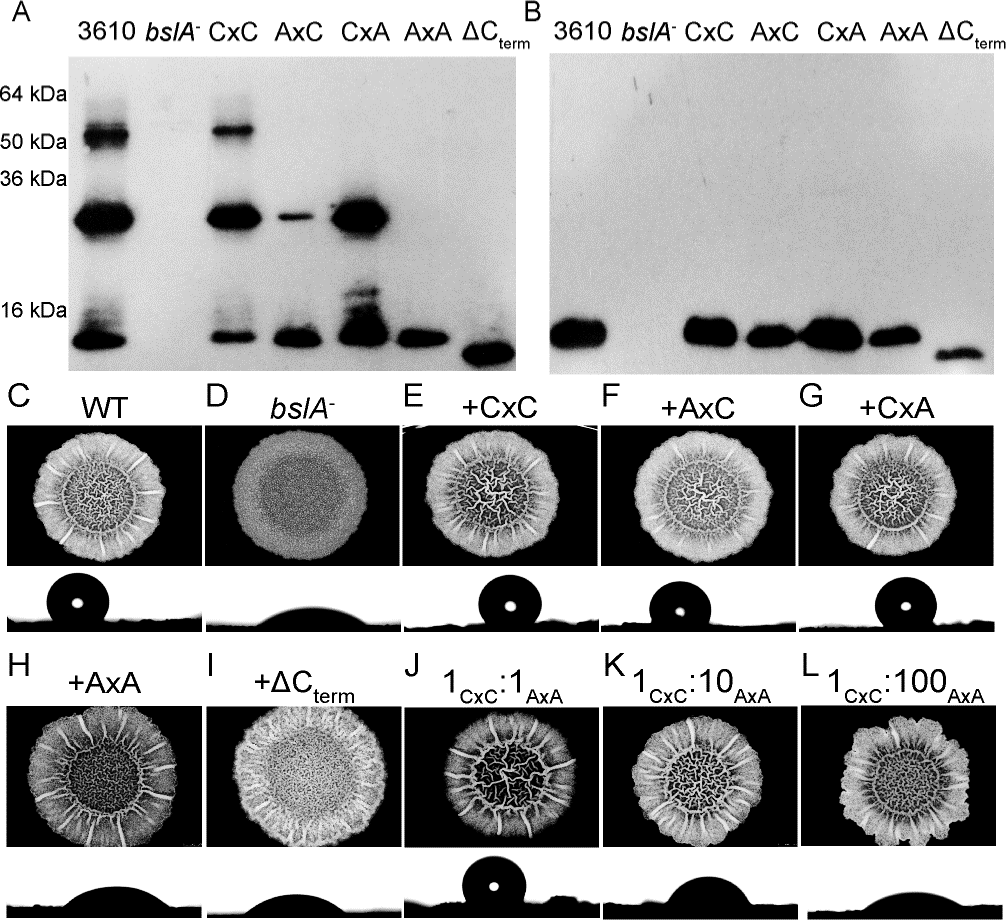
BslA is a bifunctional protein. (A, B) Western blot analysis of BslA in a native (A) and reduced (B)state using proteins extracted from biofilms (see Supplemental text S1). (C-L) Architecture and hydrophobicity of biofilms. Strains used were (C) 3610 (wild-type; NCIB3610), (D) *bslA^-^* (NRS2097), and the *bslA* mutant genetically complemented with the following variants: (E) CxC (NRS2299), (F)AxC (NR5177); (G) CxA (NRS5178), (H) AxA (NRS5179) and (I) ACterm (NRS2957). Biofilms depicted in (J, K, L) are the result of co-culturing strains NRS2299 and NRS5179. Under each biofilm is an image of a water droplet on theupper surface of the biofilm and the values of the contact angle are in Table S4.

### Genetic separation of roles in biofilm architecture and hydrophobicity

We next tested if there was a biological consequence of restricting BslA disulfide bond formation *in vivo*. Deletion of *bslA* results in a flat, featureless, wetting biofilm; our analysis reveals that production of the BsIA_C178A_, BslA_C180A_, BsIA_C178A,C180A_ and BsIA_Δ172-181_ variant proteins using the strains indicated could reinstate biofilm complexity of a *bslA* mutant to a level that was visually comparable to the wild-type strain (Fig. 1C–I). We then assessed if the biofilms formed were hydrophobic (8). As expected the wild-type strain formed a non-wetting biofilm where the water droplet had a high contact angle (124.6° ± 2.9°), whereas the *bslA* mutant was wetting (33.6° ± 2.7°) (Table S2) (8, 12). Each of the single cysteine to alanine mutations, BsIA_C178A_ (NRS5177) and BslA_C180A_ (NRS5178), displayed a non-wetting phenotype, although the contact angle calculated for the BsIA_C178A_ (NRS5177) strain was consistently lower than that measured for the wild-type strain (p <0.01; students t-test) (Table S2). In sharp contrast, the architecturally complex biofilms formed in the presence of the monomeric BslA_C178A,C180A_ (NRS5179) or BslA_Δ172-181_ (NRS2957) variants were wetting (Figure 1H and I), with contact angles that were indistinguishable from the *bslA* mutant strain (Table S2). Thus monomeric BslA seems to be associated with architectural complexity of the biofilm, whereas surface hydrophobicity requires at least the dimeric form of the protein. We note that the single point mutations, BsIA_C178A_ and BslA_C180A_, yielded only dimeric protein in addition to the monomer, (Fig 1A) indicating tetramer formation is not required. These results also reveal that the architectural complexity, which often makes a contribution to biological hydrophobicity [e.g., see (17, 18)], is not sufficient for hydrophobicity to manifest.

### Sharing BslA molecules in the biofilm

As hydrophobicity of the biofilms is at the macroscale, where BslA comprises a “common” or “public” good that can be shared by the population (19), we asked what the minimum proportion of cells producing wild type BslA was needed to achieve maximum non-wetting values. To determine this, cells expressing either the wild-type *bslA* (NRS2299) or the *bslA*_*C178A*,C180A_ (NRS5179) coding regions were co-cultured over a range of ratios, and the hydrophobicity of the resulting mature biofilm measured (Fig. 1J–L). When the strains were mixed in equal proportions, a non-wetting biofilm surface could be sustained (116.2° ± 2.8°). However, when 90% of the cells produced BslA_C178A,C180A_, the biofilm hydrophobicity remained above that of the *bslA* mutant, but there was a step-change in the wettability of the surface (50.5° ± 1.3°). Finally, wettability reached a value indistinguishable from that measured for the fully monomeric protein when 99% of the cells produced BslA_C178A,C180A_ (Table S2). These findings demonstrate that non-contributing bacteria can be tolerated during formation of the hydrophobic layer as long as they represent less than 50% of the population.

### Formation of the hydrophobic layer depends on thiol-disulfide oxidoreductases and is enhanced in an oxic environment

In the *B. subtilis* extracytoplasmic space disulfide bond formation is catalysed by one of two thiol-disulfide oxidoreductases (TDORs) named BdbA and BdbD (20–22). We predicted that if BslA was actively oligomerised then disruption of either *bdbA* or *bdbD* (or both in the case of functional redundancy) would produce a structured but wetting biofilm similar to the phenotype displayed by the monomeric BslA variants. To test this we constructed *bdbA* (NRS5552), *bdbCD* (NRS5554) (a *bdbC bdbD* operon deletion), and *bdbA bdbCD* (NRS5553) deletion strains and assessed biofilm formation and surface hydrophobicity (Fig. 2A–C) (Table S1). Consistent with our hypothesis, each of the strains formed biofilms that were morphologically indistinguishable from the parental strain NCIB3610 (Fig. 1C and Fig. 2A–C), but measurement of surface wettability revealed that the double *bdbA(C)D* mutant had a reduced contact angle of ~55° compared with the wild type strain (Table S2) (Fig. 2C). The single *bdbA* and *bdbCD* deletion strains produced surfaces with an intermediate level of hydrophobicity (Table S2) (Fig. 2A and B).

**Figure 2:**
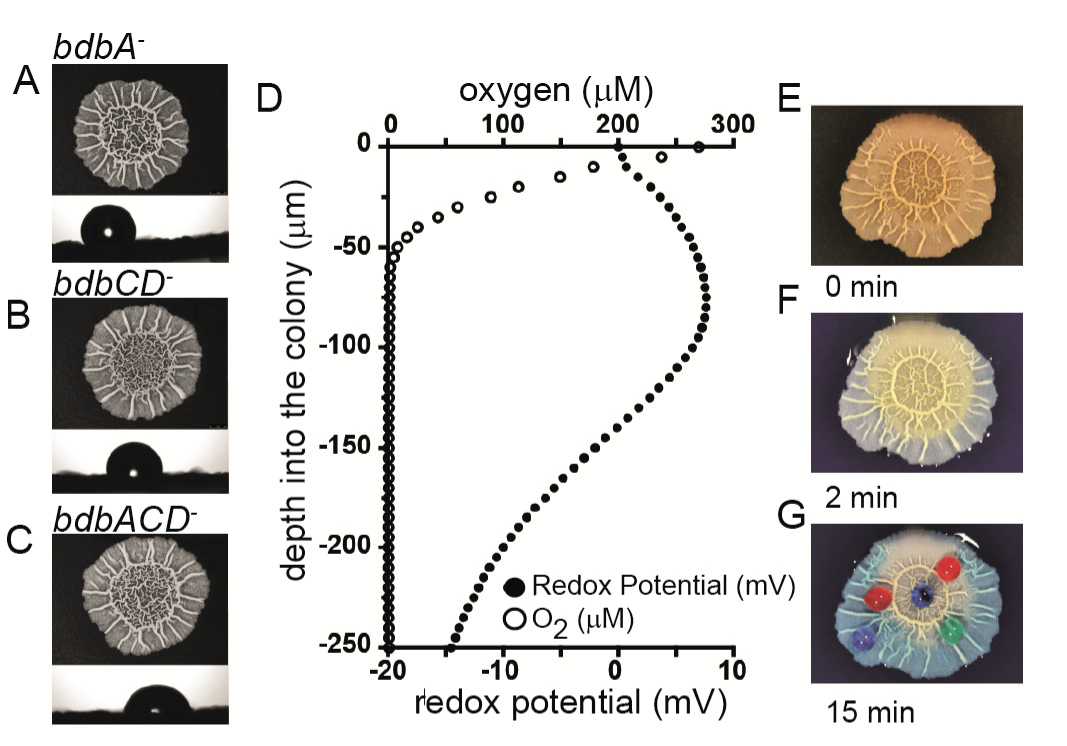
The mechanism of disulfide bond formation. Architecture and hydrophobicity of strains (A) *bdbA^-^* (NRS5552); (B) *bdbCD^-^* (NRS5554); (C) *bdbACD^-^*(NRS5553). Under each biofilm is an image of a water droplet on the upper surface of the biofilm; contact angle values are in Table S4. (D) Measurement of the oxygen concentration (μM) and redox potential (mV) as a function of the depth of the biofilm with 0 mV being set at the biofilm surface (see Supplemental textS1 for details); (E-G) Time course of water uptake by a mature biofilm visualised using pigmented water; (E) prior to treatment, 0 min; (F) 2 min after exposure; (G) 15 min after exposure. In (G) 5 ml coloured water dropletsdemonstrate retention of upper biofilm hydrophobicity.

The wettability of the *bdbA bdbCD* mutant did not reach the level measured for the *bslA* deletion strain (Table S2). Therefore, unless there is a redundant enzymatic mechanism, we propose that a combination of catalytic and spontaneous disulfide bond formation drives dimerization of BslA and leads to the formation of the hydrophobic coat. Consistent with this, measurement of the oxygen concentration through the biofilm using a microsensor demonstrated a steep decrease in the available oxygen, with no oxygen measured after 50-60 μm, as measured from the air-exposed surface, rendering the lower biofilm region anoxic (Fig. 2D). Furthermore, microelectrode-based assessment of the change in redox potential through the biofilm indicated that the upper region of the biofilm community is more oxidizing than the base (Fig. 2D). The lack of electron acceptors at the base of the biofilm suggests that BslA may be in a monomeric form in this region. Given the observation that surface hydrophobicity is associated with the ability to form dimeric protein (Fig 1C-I; Table S2), a predicted prevalence of monomeric protein at the base of the biofilm raises the possibility that this surface will be wetting. Two experimental findings support this hypothesis: first, when pigmented water is flooded onto a petri dish on which a biofilm has been grown, the dye is observed to move rapidly under the biofilm towards the centre of the community (Fig. 2E–G), indicating it is not repelled by a hydrophobic layer. Second, when the mature wild-type biofilm is turned over, such that the base is facing up, an entirely wetting surface is revealed (Table S2). Together these findings solve the conundrum (12) of how bacteria in the biofilm access nutrients, despite being surrounded by a layer of BslA.

### BslA localisation depends on disulfide bond formation

We next addressed the mechanism by which blocking oligomerization of BslA impaired the formation of a non-wetting biofilm. It is not a consequence of protein misfolding or disruption of protein production *in vivo*, as CD spectroscopy demonstrated similar secondary structures in the variant proteins (Fig. S2A) and immunoblot analysis revealed comparable levels of the BslA cysteine mutants to the wild-type protein (Fig. 1A). Therefore, we considered two non-mutually exclusive hypotheses: *1)* mutation or deletion of the C-terminal cysteine residues impairs the innate ability of BslA to form a stable elastic film that depends on lateral interactions between the monomers (12), and consequentially the ability to render the biofilm non-wetting; *2)* BslA_C178A,C180A_ cannot form a dense layer over the surface of the biofilm as observed for the wild-type protein (12). The ability of recombinant BslA_C178A_, BslA_C180A_, BslA_C178A,C180A_, and BslA_Δ172-181_ to form stable elastic protein films *in vitro* was assessed using pendant drop analysis, coupled with quantification of the wrinkle relaxation speed (12). Each of the variant proteins formed a stable protein film at the oil-water interface with no significant differences compared with wild-type BslA (Fig. S2B). Furthermore, transmission electron microscopy (TEM) indicated that the ability of the variant proteins to form an ordered 2D lattice at an interface was not impeded (Fig. S2C–E).

Next, *in situ* localisation of BslA was assessed using immunofluorescence staining of biofilm cross-sections coupled with confocal microscopy (12). The strains tested were the wild type, the *bslA* mutant, and the *bslA* mutant complemented with either the wild-type or *bslA*_C178A,C180A_ coding region. Ineach case, the strains were modified to express the *gfp* coding region to allow detection of the cells by confocal microscopy (Table S1). As has previously been observed (12), both the wild-type strain (Fig. S3A), and the *bslA* mutant complemented with the wild-type *bslA* allele, showed BslA-associated fluorescence at both the air-cell and cell-agar interfaces,consistent with formation of the BslA integument (Fig. 3A, S3C). Specificity of the immunofluorescence staining was confirmed by analysis of the *bslA* mutant (Fig. S3B).In contrast, when the *bslA* mutant was complemented with the BslA_C178A,C180A_ variant (NRS5136), BslA-linked fluorescence was not detected in high abundance at the cell-air interface and was only prevalent at the agar-cell junction (Fig. 3B, S3D). These findings indicate an inability of BslA_C178A,C180A_ either to migrate to or accumulate at the air interface, which is consistent with the wetting phenotype of the biofilm formed when this variant of BslA is produced.

**Figure 3:**
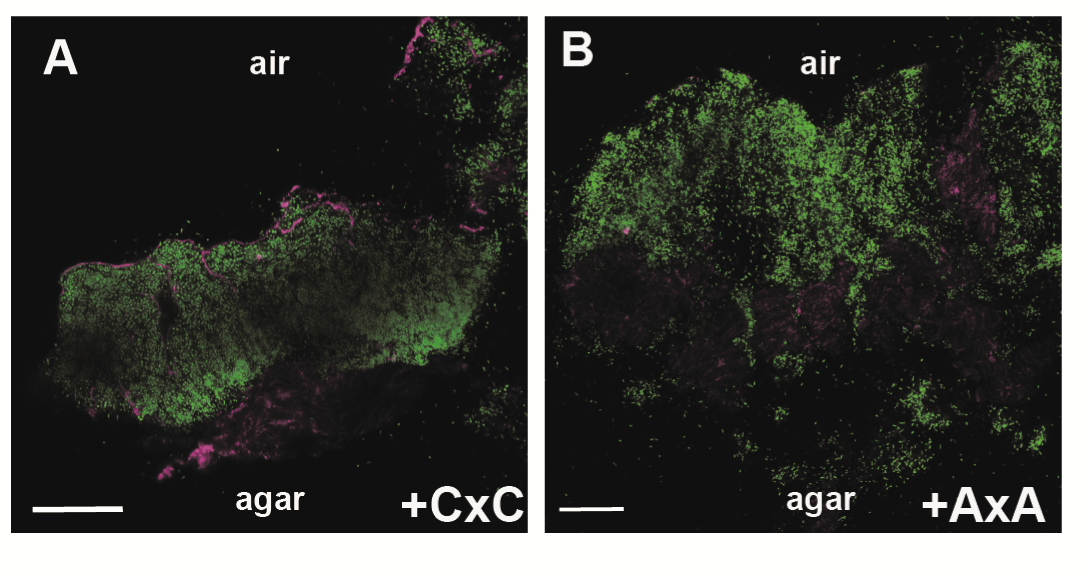
In situ analysis of BslA localisation in the biofilm. Confocal scanning laser microscopy images of biofilm cross sections through the bslA mutant complemented with either (A) the wild type variant of bslA (NRS5132); or (B) the gene encoding the BslA_C178A C180A_ variant (NRS5136) strains. Fluorescence from the GFP is shown in green and the fluorescence associated with DyLight594, representing immuno-labelled BslA staining in magenta. The scale bar represents 50 μm. The single channel data and wild-type and bslA controls are in Figure S3.

### Synthetic dimerization of the BslA paralogue

The strict requirement for dimeric BslA to mediate biofilm hydrophobicity led us to hypothesise that if the monomeric paralogue of BslA, YweA, could be engineered to dimerise *in vivo*, it may become capable of reinstating the non-wetting phenotype to the *bslA* mutant. This is not the case when the wild-type *yweA* coding region is expressed from a heterologous location in the *bslA* mutant (14). To test this, we synthesized a chimeric YweA allele (Fig. S4A), fusing the C-terminal 11 amino acids of BslA (containing C178 and C180) to the C-terminus of YweA (hereafter YweA_BslA178-181_). We additionally generated chimeric constructs where each of the cysteine residues in the BslA C-terminal region were individually, and in combination, mutated to alanine (Table S1). The wild-type YweA, chimeric YweA_BslA171-181_, and YweA_BslA171-181 C178A C180A_ recombinant proteins were purified from *E. coli* and the secondary structure assessed by CD spectroscopy. This revealed that the C-terminal extension did not significantly affect folding of the protein compared with the parental YweA protein (Fig. S5A). The oligomerisation state of the YweA_BslA171-181_ chimeric protein was assessed *in vitro* by SDS-PAGE (Fig. S4B and C) and SEC (Fig. S4D), which showed that the chimeric YweA_BslA171-181_ protein formed dimers that were reduced to the monomeric state by the addition of ß-mercaptoethanol. Bands with the apparent molecular weight expected for dimers were also formed when either one of the two cysteine residues remained (YweA_BslA171-181 C178A_ and YweA_BslA171-181 C180A_) (Fig. S4B) but only monomer was observed for the double cysteine to alanine chimeric protein(YweA_BslA171-181 C178A C180A_) (Fig. S4B). Unlike BslA, we did not detect formation of tetramers for the YweA_BslA171-181_ chimeric protein. Notably, fusion of the BslA C-terminal 11 amino acids to YweA did not alter the stability or organisation of the protein film formed *in vitro*. Pendant drop analysis, coupled with quantification of wrinkle relaxation speed, revealed that YweA, YweA_Bs1A171-181_, and YweA_Bs1A171-181 C178A C180A_ each had an average relaxation time of ~25 seconds (Fig. S5B); substantially less stable than the BslA elastic film (>600 seconds, Fig. S5B). This means that the proteins can form elastic films but they are unstable under compression. Consistent with this, TEM showed that each protein was able to form an organised lattice on a surface (Fig. S5C-E). Together, these findings indicate that we can generate YweA oligomers *in vitro* by addition of the BslA C-terminal 11 amino acids, but although they form an ordered 2D lattice, the chimeric proteins retain the fast film relaxing properties of YweA (14).

### Oligomeric YweA yields a hydrophobic biofilm

To assess the impact of engineering YweA to form intermolecular disulfide bonds *in vivo* we constructed a suite of plasmids designed to produce the YweA chimeric proteins in *B. subtilis*. The constructs were generated such that secretion through the Sec-system was directed by the BslA signal sequence(the variants are hereafter referred to as: YweA_Bs1A171-181_, YweA_BslA171-181 C178A_, YweA_BslA171-181 C180A_ and YweA_BslA171-181 C178A C180A_) (Fig. S4A). The plasmids carrying the required coding regions were introduced into the *bslA* mutant at the ectopic *amyE* locus (Table S1). Western blot analysis of proteins extracted from the biofilm, using an anti-YweA antibody, revealed that only in the presence of both cysteine residues in the appended BslA C-terminal tail could significant levels of disulfide bond-dependent oligomeric chimeric YweA protein be detected (Fig. 4A). As anticipated, the addition of a reducing agent during the extraction process rendered the protein fully monomeric (Fig. S4E). The ability of each of the YweA chimeric variants to reinstate biofilm structure and hydrophobicity to the *bslA* mutant was tested as before. As previously observed, induction of wild-type *yweA* expression did not alter the biofilm morphology or wetting phenotype, as comparedwith the parental *bslA* mutant (Fig. 4B) (14). In sharp contrast, appending the BslA C-terminal 11 amino acids to YweA allowed the chimeric YweA_BslA171-181_ protein to return hydrophobicity to the *bslA* mutant strain (114.8° ± 1.1°) (Fig. 4B) (Table S2). It should however be noted that the biofilm structure retained more architectural similarities with the *bslA* mutant, rather than the wild-type NCIB3610 strain (Fig. 4B). Expression of YweA_BslA171-181_ thus results in an unstructured, yet non-wetting, phenotype. The slightly lower contact angle measured (114.8° ± 1.1° *vs* 124.6° ± 2.9° for the structured, wild-type biofilms) may indicate that biofilm architecture makes some contribution to overall hydrophobicity, as has been suggested elsewhere (18). In contrast, when the C-terminal extension was mutated such that one (NRS5515 and NRS5516), or both (NRS5210) cysteine residues were replaced with alanine, yielding monomeric BslA *in vivo*, the colonies formed retained both the wetting and unstructured phenotypes exhibited by the *bslA* mutant (compare Fig. 1D with Fig. 4B).Thus, dimeric chimeric YweA containing the two cysteine residues is able to reinstate biofilm hydrophobicity, whereas monomeric chimeric YweA is unable to reinstate biofilm architectural complexity.

### Hydrophobicity and architecture play a role in resistance to chemical attack

The ability to genetically separate biofilm structure from hydrophobicity allowed us to assess if protection to external insult is conferred to the bacteria by blocking access of the chemical and / or mediated by the architectural complexity. *B. subtilis* biofilms are non-wetting when challenged with selected commercial biocides (6) with chlorhexidine gluconate being the reactive agent of a number of such biocides. Analysis reveals that 1% (v/v) chlorhexidine gluconate is non-wetting on “wild type” biofilms (Fig. 5A) and in contrast is wetting on both the *bslA* mutant and the variant carrying the monomeric BslA_C178A,C180A_(Fig. 5A, Table S2), providing an ideal test agent to test the question posed above. The number of surviving cells after treatment for 5 minutes with 1% (v/v) chlorhexidine gluconate was calculated relative to a comparable sample exposed to saline solution. We noted a clear role for BslA in mediating cell survival: in the absence of *bslA*, exposure to 1% (v/v) chlorhexidine gluconate resulted in ~38-fold fewer surviving cells compared with when BslA was produced (Fig. 5B). In contrast, when BslA_C178A,C180A_ was present, resulting in a structured but wetting biofilm, limited cell survival was measured. Cell survival was ~5-fold lower than that measured for the “wild type” strain (Fig. 5B). As BslA is known to have a role in sporulation specifically during biofilm formation (19), we calculated the level of heat resistant spores in each biofilm at the time point used to test chemical resistance. As established previously the level of sporulation was low in the *bslA* mutant (Fig. 5C), however the number of spores formed in the presence of BslA_C178A,C180A_ was the same as when wild type BslA was produced (Fig. 5C). *In toto* these data demonstrate that the protection received by the cells in the biofilm results from a combination of the hydrophobicity and structure conferred by BslA, a phenomenon only revealed by the ability to genetically separate the two functional contributions.

**Figure 5:**
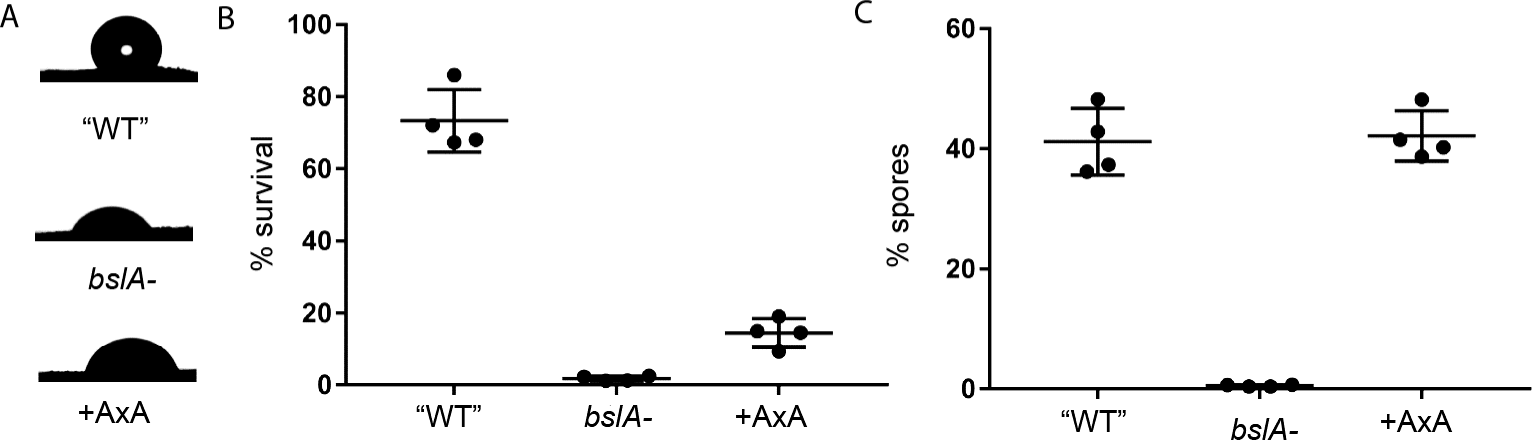
Cell survival after exposure to chlorhexidine gluconate. (A) An image of a 1% (v/v) chlorhexidine gluconate droplet on the upper surface of the biofilm for strain “WT” (NRS5132); *bslA* mutant (NRS5131), and the *bslA* mutant complemented with BslA_C178A C180A_ “+AxA” (NRS5136); the values of the contact angle are in Table S4; (B) The strains described above were exposed to 1% (v/v) chlorhexidine gluconate and the percentage survival calculated; (C) Percentage sporulation was calculated for the strains detailed in (A). The error bars represent the standard deviationfrom the mean.

## Discussion

The mechanism revealed here for controlling protein function through oxidation allows us to classify BslA as a bifunctional protein with genetically separable roles in biofilm formation, mediated by cysteine residues in the C-terminal tail. BslA is a key component of the *B. subtilis* biofilm and plays a role in both biofilm architecture and hydrophobicity (8, 12, 23). Dimerization of BslA directly facilitates the development of the non-wetting layer and relies on a cysteine motif (CxC) at the C-terminus. Monomeric variants of BslA yield biofilms that are architecturally complex but lack hydrophobicity, while conversely, transplantation of the BslA C-terminus onto the normally monomeric non-functional paralogue YweA allows the chimeric protein to dimerise and consequentially rehabilitate hydrophobicity, but not architecture, to a *bslA* mutant. These findings are consistent with the accumulation of BslA at the air-biofilm interface only when it can dimerize, as observed by high resolution *in situ* immunofluorescence microscopy (Fig. 3). The BslA layer is thus shared by the entire community, and can form over and shield non-producing bacteria (Fig. 1J–L). By virtue of being able to separate the role of BslA in biofilm architecture from hydrophobicity we have shown that both architectural complexity and the hydrophobic layer contribute to protecting the residents from biocides, thus providing an evolutionary advantage to the resident cells. Finally, we have demonstrated that an effective hydrophobic barrier can be generated by a subpopulation of the biofilm and this finding raises the question of why the whole population has evolved to produce BslA (12) when other biofilm matrix molecules are produced in a bimodal manner (24).

A model can be postulated that explains formation of a BslA layer that is more than one molecule (or dimer) deep at the air interface (Fig. 6). It is known that BslA dimers are orientated longitudinally (tail-to-tail) with only one cap of each dimer able to interact with the air interface at a time (11). The orientation of tetrameric forms of the protein is unclear, however as dimeric protein is sufficient to give rise to the observed effects, it is not considered further here. Based on the premise that the hydrophobic cap of one dimer can serve as a hydrophobic interface for another dimer, it leads to a scenario where the integument of the biofilm could be comprised of the dimeric proteins layered against each other (Fig. 6). Furthermore, as monomeric and dimeric BslA are able to coexist *in vitro* in a protein film, the mature *in vivo* hydrophobic layer has the potential to be generated from a mixture of monomeric and dimeric protein. We have confirmed that stable *in vitro* film formation is not needed to confer hydrophobicity; the chimeric YweA_BslA 171-181_ variant is able to render the community hydrophobic but forms a protein film that relaxes quickly under compression (Fig. 4B and S5B). This is consistent with recent analysis of BslA orthologues where *B. pumilus* variant lacked stability in the elastic film but could substitute for the *B. subtilis bslA* gene *in vivo* (14). Together these data indicate that stability in the lateral interactions between the BslA molecules is independent of the ability to confer hydrophobicity to the community but may determine architectural complexity. However, details of the lateral interaction sites between the protein molecules are currently unknown.

**Figure 6:**
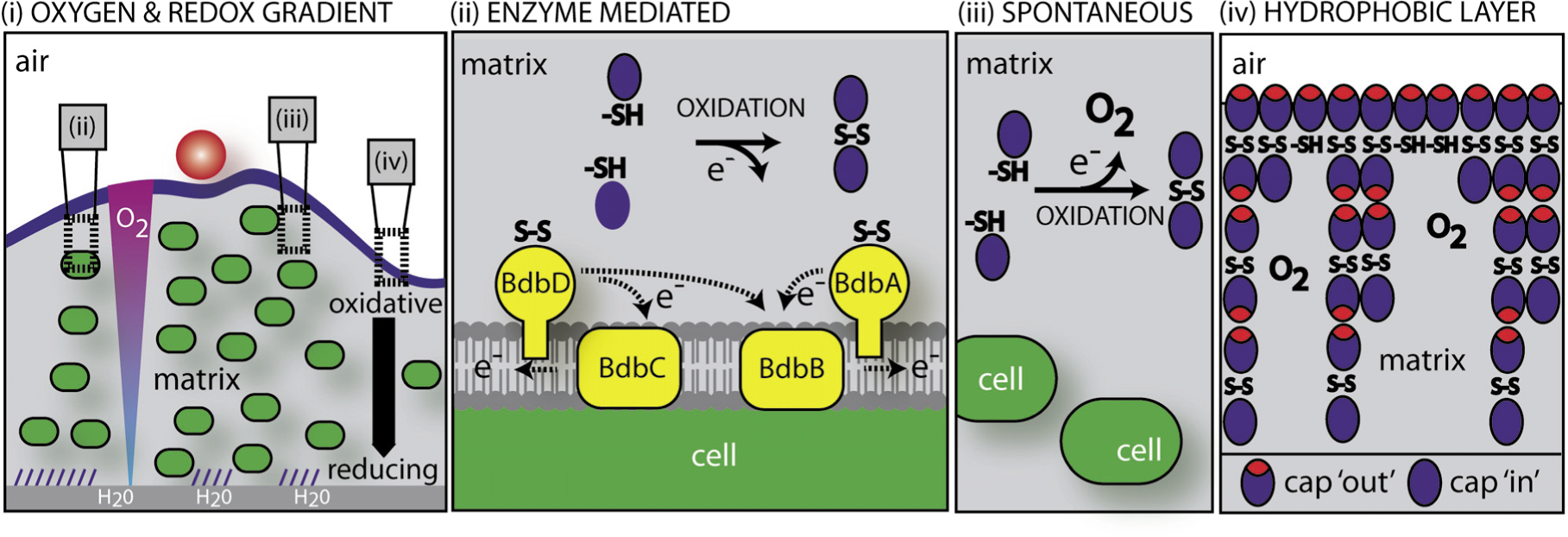
Model of BslA function. (i) Schematic of a biofilm cross-section depicting the oxygen and redox gradient through the depth of the structure. The blue layer at the air interface represents the BslA hydrophobic coat with a water droplet on top (red circle) and the hatched lines at the base represent BslA in a form that allows water and nutrient uptake into the community. The bacteria are green ovals and the agar surface is the grey zone at the base of the biofilm; (ii) Oligomerisation of BslA is mediated by thiol-oxidoreductases that reside in the membrane and extracytoplasmic space. The electrons (e) released by oxidation upon formation of the disulfidebond flow into the respiratory chain. BslA is shown as a blue oval in the biofilm matrix (grey). For simplicity only one disulfide bond has been depicted (S-S). The reduced form of the protein is represented by the (-SH) annotation; (iii) Oligomerisation of BslA is also likely to occur using molecular oxygen as the electron acceptor; (iv) depicted is a model for how the BslA coat might present in the biofilm as a mixture of oligomers. The ability of BslA to present in a cap ‘in’ and cap ‘out’ configuration is represented wherethe cap ‘out’ form is adopted by the molecules at a hydrophobic interface.

**Figure 4:**
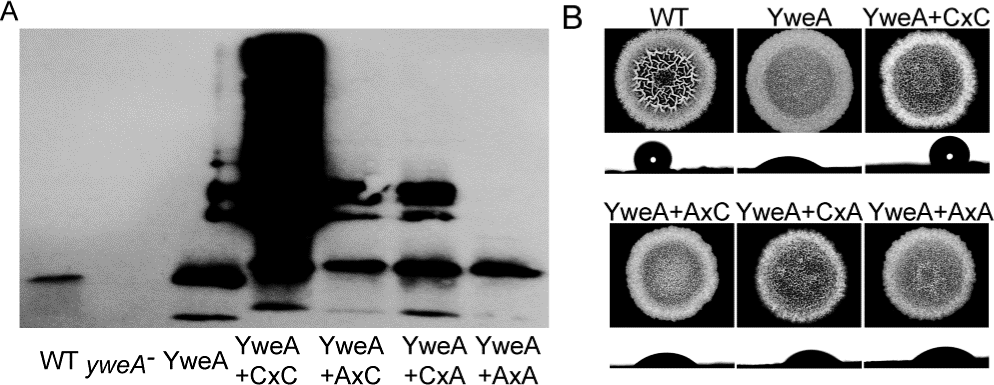
Forcing dimerisation of YweA can rehabilitate biofilm hydrophobicity in a *bslA* mutant.Western blot analysis of YweA in a native (A) state detected from biofilm protein extracts (see Supplemental text S1). Strains used were 3610 (WT; NCIB3610), *yweA^-^* (NRS2405), and the *bslA*mutant engineered to produce the following YweA variants: YweA (NRS5551), YweA_BslA171-181_ (NRS4834); YweA_BslA171-181 C178A_ (NRS5515) YweA_BslA171-181 C180A_ (NRS5516)and YweA_BslA171-181 C178A C180A_ (NRS5210). (B) Biofilms using strains detailed in part (A); Under each biofilm is an image of a water droplet onthe upper surface of the biofilm: values of the contact angle are in TableS4 and strains as part (A).

BslA is therefore multifunctional across three different axes: *1)* dimerization (and/or tetramerisation) contributes to hydrophobicity, whereas monomeric protein is sufficient for biofilm complex architecture; *2)* the “cap out” form of the protein renders a surface layer hydrophobic whereas we can infer that a “cap in” form is present in the wetting protein layer at the base of the biofilm where the protein is exposed to water (11, 13); and *3)* stable lateral interactions between BslA molecules, which can be measured *in vitro*, appear to be required for architectural complexity, whereas unstable lateral interactions (reflected in unstable film formation *in vitro)* nonetheless are sufficient to give rise to hydrophobicity if dimeric protein is present. We have previously elucidated a link between the behaviour of the cap region and the strength of lateral interactions (13), such that these two functionalities appear to influence each other. Conversely, the behaviour of the cap region of BslA is functionally independent from the protein oligomerisation state, as both monomeric protein and dimeric protein are able to flip between ‘cap in’ and ‘cap out’ conformations. Thus, the bottom of the biofilm is wetting not just because the protein is monomeric but because the cap region is also exposed to water; likewise, the top of the biofilm is hydrophobic not just because BslA forms dimers and tetramers but because the cap region is also exposed to air. The mechanisms by which monomeric BslA gives rise to an architecturally complex biofilm, and the reasons why dimeric protein (at least) is required to give rise to a hydrophobic coat, remain to be elucidated, but possibly involve (an) interaction(s) with other components of the biofilm.

## Bifunctionality through dimerisation

Cysteine residues, possessing a thiol group (15), have specialized roles in a wide range of cellular processes and control protein folding and stability, multimerisation, and function through disulfide bond formation (25). It is clear from our secondary structural analysis (Fig. S2) that the role for disulfide bond formation in modulating BslA activity is not linked with either protein stability or folding, but is instead associated with imparting new function through multimerisation. This is in contrast to the fungal hydrophobins where disulfide bonds stabilize the structure of the protein (26) and is more analogous to the mechanism used to control activity of the von Willebrand factor (vWF) during blood clotting (25). Previous bioinformatic analyses reveal that Firmicutes, including *B. subtilis*, limit the number of proteins that contain cysteine residues (27). Consistent with this there are no essential components that require disulfide bond formation in the *B. subtilis* biofilm, as deletion of the known extracytoplasmic thiol oxidoreductases, *bdbA* and *bdbD* does not lead to pleiotropic defects. It is only when the integrity of hydrophobicity is assessed that differences in the surface wettability are uncovered (Table S2). It has been proposed that in Firmicutes proteins that contain disulfide bonds have roles in “niche” functions which are not linked with essential cellular processes: for example genetic competence that allows the uptake of exogenous DNA (21, 28) and production of the S-linked glycopeptide sublancin that has antimicrobial properties (23, 29, 30).Arguably, biofilm hydrophobicity would fit within this category, and consistent with genetic competence and sublancin production, successful formation of the hydrophobic layer imparts a fitness advantage by, in this case, excluding chemicals from entering the biofilm.

## Heterogeneity in electron acceptor availability drives BslA function

In addition to catalysis-driven disulfide bond formation by extracytoplasmic thiol-oxidoreductases, spontaneous disulfide bond formation using molecular oxygen (or via a redox active molecule) as the direct electron acceptor enhances BslA dimerization at the air-biofilm surface interface. This is driven in the top layer of the biofilm by an oxidative, oxygen rich zone that is conducive to disulfide bond formation (Fig. 6) (31). It is well established that oxygen gradients stratify natural, mixed species, biofilms by driving distribution of microorganisms with different respiratory requirements (32–34). Furthermore, the availability of electron acceptors, including oxygen, can drive structuring of biofilm morphology, as has been shown for *Pseudomonas aeruginosa* (35, 36). Therefore, these findings expand the role that oxygen gradients can play in biofilm formation by highlighting an ability to modulate protein function. The ability to respond to localised oxygen heterogeneity in this manner provides an efficient mechanism for bacteria to maximise resource utilisation by generating bifunctionality through redox sensitivity. Furthermore, BslA dimerization in response to the oxygen gradient solves the conundrum of how *B. subtilis* obtains nutrients when it is surrounded by a layer of BslA (12); i.e. the anoxic base of the biofilm is hydrophilic despite the abundance of BslA (Fig. 6). We predict that bifunctionality of matrix components will emerge as a common theme across the species. Consistent with this, the *Vibrio cholerae* biofilm matrix protein, RmbA, is proteolytically cleaved to a form that promotes recruitment of cells to the biofilm in an exopolysaccharide independent manner; effectively the proteolytic event changes the function of RmbA (37). Additionally, flagella synthesized by *Escherichia coli* are used, not only for motility, but for imparting structural rigidity to the community by entwining the bacterial cells (38). Control of bifunctionality for each of these examples is distinct and raises the question of how many different mechanisms have evolved to maximise use of a limited number of components.

## Materials and methods

### General growth conditions and strain construction

The *Bacillus subtilis* and *Escherichia coli* strains used and constructed in this study are detailed in Table S1. *E. coli* strain MC1061 was used for the construction and maintenance of plasmids. *B. subtilis* 168 derivatives were obtained by transformation of competent cells with plasmids using standard protocols (39). SPP1 phage transductions were used to introduce DNA into *B. subtilis* strain NCIB3610 (40). Both *E. coli* and *B. subtilis* strains were routinely grown in Lysogeny-Broth (LB) medium (10 g NaCl, 5 g yeast extract, and 10 g tryptone per litre) at 37°C for 16 hours. For complex colony formation *B. subtilis* strains were grown on MSgg medium (5 mM potassium phosphate and 100 mM MOPS at pH 7.0 supplemented with 2 mM MgCl_2_, 700 μM CaCl_2_, 50 μM MnCl_2_, 50 μM FeCl_3_, 1 μM ZnCl_2_, 2 μM thiamine, 0.5% glycerol, 0.5% glutamate) (5) solidified with 1.5% Select Agar (Invitrogen) at 30 °C for 48 hours (5) and imaged as described previously (40). Ectopic gene expression was induced by medium supplementation with 25 μM isopropyl β-D-1-thiogalactopyranoside (IPTG) as indicated. When appropriate, antibiotics were used at the following concentrations: ampicillin 100 μg ml^-1^, chloramphenicol 5 μg ml^-1^, erythromycin 1 μg ml^-1^ with lincomycin 25 μg ml^-1^, kanamycin 25 μg ml^-1^, and spectinomycin 100 μg ml^-1^. *AbdbA, AbdbCD* and *AbdbACD* mutant strains were constructed as described in Table S1 using standard methodologies.

### Plasmid construction and site-directed mutagenesis

All strains, plasmids and primers used in this study and presented in Tables S1, S3 and S4 and were constructed using standard methods. The plasmids for BslA_42-181_ overproduction were obtained by site-directed mutagenesis using the plasmid pNW1128 as template, which is a pGEX-6P-1 derivative used previously to overexpress BslA_42-181_ (41). Primers for the codon substitutions are included in Table S4 and mutagenesis was achieved following Stratagene Quikchange kit recommendations. The plasmids for overexpression *yweA* or the fusion *yweA-BslA_171-181_* were obtained using standard techniques with either NCIB3610 or a synthetic DNA construct as the template DNA during PCR. The sequence of the synthetic construct generated by Genescript was as follows: 5’-agatct CGA TCT gca tca atc gag gca aaa aca gtt aac agc acg aaa gag tgg acc att tct gat att gaa gtg aca tat aaa cca aat gcg gtg ctt tct ctt gga gcg gta gaa ttt caa ttt cct gac ggg ttt cat gct acg aca aga gat tca gtg aat gga aga aca ctg aaa gaa aca cag att tta aac gat gga aaa aca gtc aga ctc ccg ctt acg ctt gat ttg tta ggc gca tcc gaa ttt gac ctt gtc atg gtg cgt aaa act ctt cct cgc gca ggc act tac acg att aaa ggc gat gta gta aac ggt ttg gga atc ggc agt ttt tat gct gaa acg cag ctg gtg att gat ccc cgt **agc act cct ccg act cag cct tgc ggt tgc aac taa** GCA TGC-3’. The sequence in capital letters represents restriction sites, the sequence in bold represents the C-terminal *bslA* coding region and finally the sequence in lower case represents the mature region of the *yweA* gene sequence.

### Biofilm hydrophobicity contact angle measurements

Biofilm hydrophobicity was evaluated by measuring the contact angle of a 5 μl droplet of water or 1% (v/v) chlorhexidine gluconate placed on the upper or lower surface of the biofilm that had been grown for 48 hours at 30°C. Measurements were obtained using a ThetaLite TL100 optical tensiometer (Biolin Scientific) and analysed with OneAttension. The water droplet was allowed to equilibrate for 5 minutes prior imaging and measurement. Contact angles are present as the average of at least 3 independent experiments and the standard error of the mean associated with these values (Table S2).

### Protein Purification

BslA4_1-181_ protein and its derivatives were over expressed and purified as previously described (12). Briefly, the pGEX-1-6P derivative plasmids were introduced into *E. coli* BL21 (DE3). After growth and overexpression, the protein was purified in HEPES buffer. To achieve this, the *E. coli* cells were lysed in an Emulsiflex cell disruptor and the solubilized protein extracts, cleared of cell debris, were incubated with Glutathione Sepharose 4B (GE Healthcare), allowing the fused protein to bind to the GST binding beads. After incubation, the beads containing the BslA-fusion were recovered using a gravity flow column (Bio-Rad) and suspended into new buffer containing DTT and TEV-His tagged protease. TEV removes the GST tag from BslA protein, which remains soluble in the purification buffer. The protease and unbound GST are then separated from BslA by incubating the mixture with Ni-nitrilotriacetic acid agarose (Qiagen) and Glutathione beads, followed by a new passage in a gravity flow column. The flow-through recovered contains the purified protein, which is then concentrated using VivaSpin concentrator. Protein quality was confirmed by size exclusion chromatography (SEC) analysis.

To purify YweA_31-155_ and the chimeric proteins YweA-BslA_171-181_ and its derivatives YweA-BslA_171-181 C178A_, YweA-BslA_171-181 C180A_ and YweA-BslA_171-181 C178A C180A_ the required coding regions were introduced into pET-15b-TEV vector (see Tables S3 and S4). The pET-15b-TEV derivative plasmids were introduced into *E. coli* BL21 (DE3). After growth and overexpression *E. coli* cells were suspended in HEPES buffer (300 mM NaCl, 50 mM HEPES pH8 with the addition of 20 mM imidazole) containing protease inhibitors (Roche complete EDTA-free) and lysed using the Emulsiflex C3. The solubilized protein extracts, cleared of cell debris, were incubated with Ni-nitrilotriacetic acid agarose(Qiagen) for 4 hours, allowing the fused protein to bind to the Ni-NTA binding beads. After incubation, the beads containing the YweA-fusion proteins were recovered using a gravity flow column (Bio-Rad) and suspended into new buffer (300 mM NaCl, 50 mM HEPES pH 8.0 with the addition of 250 mM imidazole) on column for half an hour to promote the protein release. Proteins were recovered by collection of the column flow-through which was incubated overnight in buffer 300 mM NaCl, 50 mM HEPES pH 8.0 with the addition of DTT and TEV-His tagged protease. VivaSpin concentrators were used to remove the imidazole in solution by replacement up with the purification buffer. The protein recovered from the concentrators was incubated overnight with Ni-NTA beads to bind to the His-tag fragments released, followed by a new passage in a gravity flow column. The flow-through recovered contains the purified protein, which is then concentrated using VivaSpin concentrators. Protein quality was confirmed by size exclusion chromatography (SEC) analysis.

### Size exclusion chromatography (SEC)

To evaluate the presence or absence of dimers in the purified proteins, 500 ng of purified protein was analyzed by size exclusion chromatography using a Superdex 75 10/300GL column with a low rate of 0.5ml/min in the original purification buffer. Protein samples were also analyzed using a 14% SDS-PAGE, with and without the addition of a reducing agent (β-mercaptoethanol) as the loading dye and stained with Instant Blue prior to photography.

### Circular Dichroism (CD) analysis

CD analysis was performed using a Jasco J-810 spectropolarimeter. Samples were measured in a 0.1 cm quartz cuvette at 0.15 mg/ml protein concentrations in 25 mM, pH7 phosphate buffer. All samples were measured using a 50 nm/s scan rate, 0.1 nm data pitch and a digital integration time of 50 nm/s. Twenty individual spectra were measured and averaged per sample.

### Transmission Electron Microscopy (TEM)

A 5 μl droplet of 0.025 mg/ml protein solution (in pH7, 25 mM phosphate buffer) was pipetted onto a carbon-coated copper grid (TAAB Laboratories Equipment Ltd) and left for 5 minutes before being wicked away from the side using filter paper. Subsequently, a 5 ul droplet of 2% uranyl acetate was pipetted onto the grid. This was also left for 5 minutes before being wicked away from the side. Stained grids were then imaged using a Philips/FEI CM120 BioTwin transmission electron microscope.

### Wrinkle relaxation

0.2 mg/ml protein samples (in pH7, 25 mM phosphate buffer) were loaded into a glass syringe with a needle diameter of 1.83 mm. A 40 μl droplet of protein solution was expelled into glycerol trioctanoate oil and allowed to equilibrate for 20 minutes at room temperature. Subsequently, the droplet was compressed by retraction of 7 μl from the droplet, resulting in the formation of wrinkles in the protein film. Images of the droplet are taken from the moment of compression using a CCD camera. Images were acquired at a rate of 30 frames per second (fast relaxers-all YweA variants) or 2 frames per second (slow relaxers-all BslA variants) for up to 10 minutes. Relaxation of the wrinkles was analyzed in ImageJ. A line profile was drawn across the wrinkles and greyscale values (0-255) were extracted for every pixel along the line. Greyscale values were background corrected and normalized. At least five independent experiments were performed per protein.

### Oxidative cross-linking of cysteines in colony biofilms

*B. subtilis* 48 hour grown complex colonies were collected from the agar plate using a sterile loop and suspended in 250 μL of LB. The biomass was disrupted by passage through a 23 X 1 needle 10 times. The cell suspension obtained was incubated for 15 min at 37°C with 1.8 mM Cu(II)-(o-phenanthroline)_3_ (hereafter CuPhe) or 10 mM Dithiothreitol (DTT), followed by centrifugation and wash of the pellet with 250μl PBS. After centrifugation the pellet was suspended in 250 μl PBS followed by a 15 minute incubation with 8 mM N-ethylmaleimide/10 mM EDTA to stop the reaction (42, 43). Samples were then centrifuged and the pellet washed with 250 μl PBS. The cell pellet was suspended in 250 μl of BugBuster Master Mix (Novagen) followed by gentle sonication to promote the release of the proteins from the biofilm matrix. The samples were then incubated at room temperature with agitation for 20 min, and the insoluble cell debris was removed by centrifugation at 17,000 × g for 10 min at 4 °C. The proteins samples were then analysed by Western blot.

### Western Blot analysis

1.5 μg of total protein extract (see above) was separated on a14 % SDS-PAGE before transfer onto PVDF membrane (Millipore) by electroblotting at 25 V for 2 hours. The membrane was incubated for 16 hours in 3% (wt/vol) powdered milk in TBS [20 mM Tris·HCl (pH 8.0) and 0.15 M NaCl] at 4°C with shaking. This was followed by 2 hours incubation with purified anti-BslA antibody at a dilution of 1:500 (v/v) inTBS in 3% powdered milk wash buffer (TBS + 0.05% Tween 20). The membrane was washed using wash buffer (TBS + 0.05% Tween 20) and incubated for 45 min with the secondary antibody conjugated to horseradish peroxidize [goat anti-rabbit (Pierce)] at a dilution of 1:5,000. The membrane was washed, developed, and exposed to X-ray film. Western blot analysis for YweA detection was performed as above with purified anti-YweA antibody at a dilution of 1:10,000 (v/v).

Oxygen profiling of biofilms. Overnight cultures of *Bacillus subtilis* strain 3610 were inoculated from a streaked plate and subsequently grown in lysogeny broth (LB) at 37°C with shaking at 250 rpm for 12-16 hours. Precultures were diluted ten-fold in LB and grown at 37°C with shaking at 250 rpm until OD_600nm_ ≈ 1.0. Five microliters of the culture were spotted onto an MSgg agar plate and grown at 30°C for two days prior to analysis. A 25 μm-tip Clark-type oxygen microsensor (Unisense OX-25)was used to measure the oxygen concentrations. The oxygen microsensor was calibrated according to manufacturer’s instructions and measurements were taken throughout the depth of the biofilm (step size = 5 μm, measurement period = 3 seconds, wait time between measurements = 3 seconds). Four different colonieswereprobed, and representative data are shown.

### Redox profiling of biofilms

Biofilms were grown as above and a 25 μm-tip redox microelectrode with an external reference (Unisense RD-25 and REF-RM) was used to measure the extracellular redox potential. After calibrating the redox microelectrode according to manufacturer’s instructions, redox measurements were taken throughout the depth of the biofilm (step size = 5 μm, measurement period = 3 seconds, wait time between measurements = 5 seconds). The redox potential was set to zero at the surface of the colony and relative values are plotted. Three different colonies were probed, and representative data are shown.

### Sporulation Assay

For heat resistant spore quantification, colony biofilms were grown for 48h at 30°C. Cells were collected in 1 ml of saline solution, disrupted by passage through a 23 X 1 needle 10 times and subsequently subjected to mild sonication (20% amplitude, 1 second on, 1 second off for 5 seconds total) to liberate bacterial cells from the matrix. To kill vegetative cells, the samples were incubated for 20 min at 80°C. To determine viable cell counts, serial dilutions were plated before and after the 80°C incubation on LB agar supplemented with 100 μg ml^-1^ spectinomycin. The percentage of spores was established by colony forming unit counting and results are presented as the percentage of colony forming units obtained after incubation of the samples for 20 min. at 80°C divided by the number of colony forming units obtained before the heat inactivation.

### Cell survival upon chlorhexidine gluconate exposure

To test biofilm resistance to chlorhexidine gluconate, colony biofilms were grown for 48 hours at 30°C. Droplets of 5 μl of 1% (v/v) chlorhexidine gluconate were placed on the biofilm surface (near the periphery) for 5 minutes at room temperature. The solution was removed and a punch biopsy of 5 mm in diameter recovered (this area encompassed the entire exposed region). This sample was transferred immediately into 500 μl of saline solution this diluting any remaining chlorhexidine gluconate, disrupted by passage through a 23 X 1 needle 10 times and washed once with saline solution. The sample was subsequently subjected to mild sonication (20% amplitude, 1 second on, 1 second off for 5 seconds total) to liberate bacterial cells from the matrix. Serial dilution of the cell suspension were made and 100 μl plated onto LB agar supplemented with 100 μg ml^-1^ spectinomycin; saline solution was used as a control for the process. The percentage survival was evaluated by colony forming unit counting and results are presented as the percentage of colony forming units obtained after exposure to 1% (v/v) chlorhexidine gluconate divided by the number of colony forming units recovered on the control spots.

### Immunofluorescence

To prepare cross sections for microscopy, biofilms were grown from strains constitutively expressing GFP were grown for 2 days in standard conditions (see Table S1) as previously described (12). A quarter of the colony was excised and placed into optimum cutting temperature compound (OCT, AgarScientific) and frozen in iso-pentene chilled with liquid nitrogen. A Leica CM3050 S cryomicrotome was used to cut cross sections of this colonies (10–12 μm), which were placed onto SuperFrost Ultra Plus adhesion microscope slides (VWR). The cross sections were then fixed for 10 minutes in para-formaldehyde (4% in TBS), washed 3 times with TBS and blocked overnight in 2% (w/v) fish gelatine (Sigma) prepared in TBS. After washing the cross sections, anti-BslA antibody diluted 1:200 in AbDil solution (2% BSA, 0.1% azide in TBS) was applied into the cross sections and incubated for 2.5 hours at room temperature. After 3 washes of 5 minutes each with TBS, the secondary antibody was applied in a dilution of 1:150 in AbDil and incubated for 90min at room temperature. The secondary antibody used was DyLight594-conjugated Affinity Pure Donkey Anti-Rabbit IgG (H+L) (Jackson ImmunoResearch). After the incubation, the coverslips were washed 3 times with TBS (5 minutes each) and mounted in anti-fade containing medium [0.5% p-Phenylenediamine (Free base; Sigma), 20 mM Tris (pH 8.8), and 90% glycerol] [Cramer, L., and Desai, A. (1995) (http://mitchison.med.harvard.edu/protocols/gen1.html)] and sealed with nail varnish. Samples were imaged using a Zeiss LSM700 confocal scanning laser microscope fitted with 488-nm and 555-nm lasers and an EC PlanNeofluar 40×/ NA 1.30 oil differential interference contrast (DIC) M27 or an alpha Plan-Apochromat 100×/ NA 1.49 oil DIC objective. Images were captured and analysed using Zen2011software, which was also used to prepare the images presented.

## Acknowledgements

Work was supported by the Biotechnology and Biological Sciences Research Council [BB/L006804/1; BB/L006979/1; BB/M013774/1; BB/N022254/1]. We acknowledge the Dundee Imaging Facility, Dundee, supported by the ‘Wellcome Trust Technology Platform’ award [097945/B/11/Z] for help with experiments. JJ was supported by NIH training grant 5T32GM008798, LEPD was supported by NIH grant R01AI103369. We are very grateful to Drs. L Hobley and L Cairns for initial observations, Prof. F Sargent for helpful discussions, Dr. A Ostrowski and Ms. E. Bissett for plasmids and strains, and Profs. Ben-Yehuda and van Dijl for the anti-YweA antibody and the *bdbDC* mutant respectively.

## Suplementary-material

**Figure S1:**
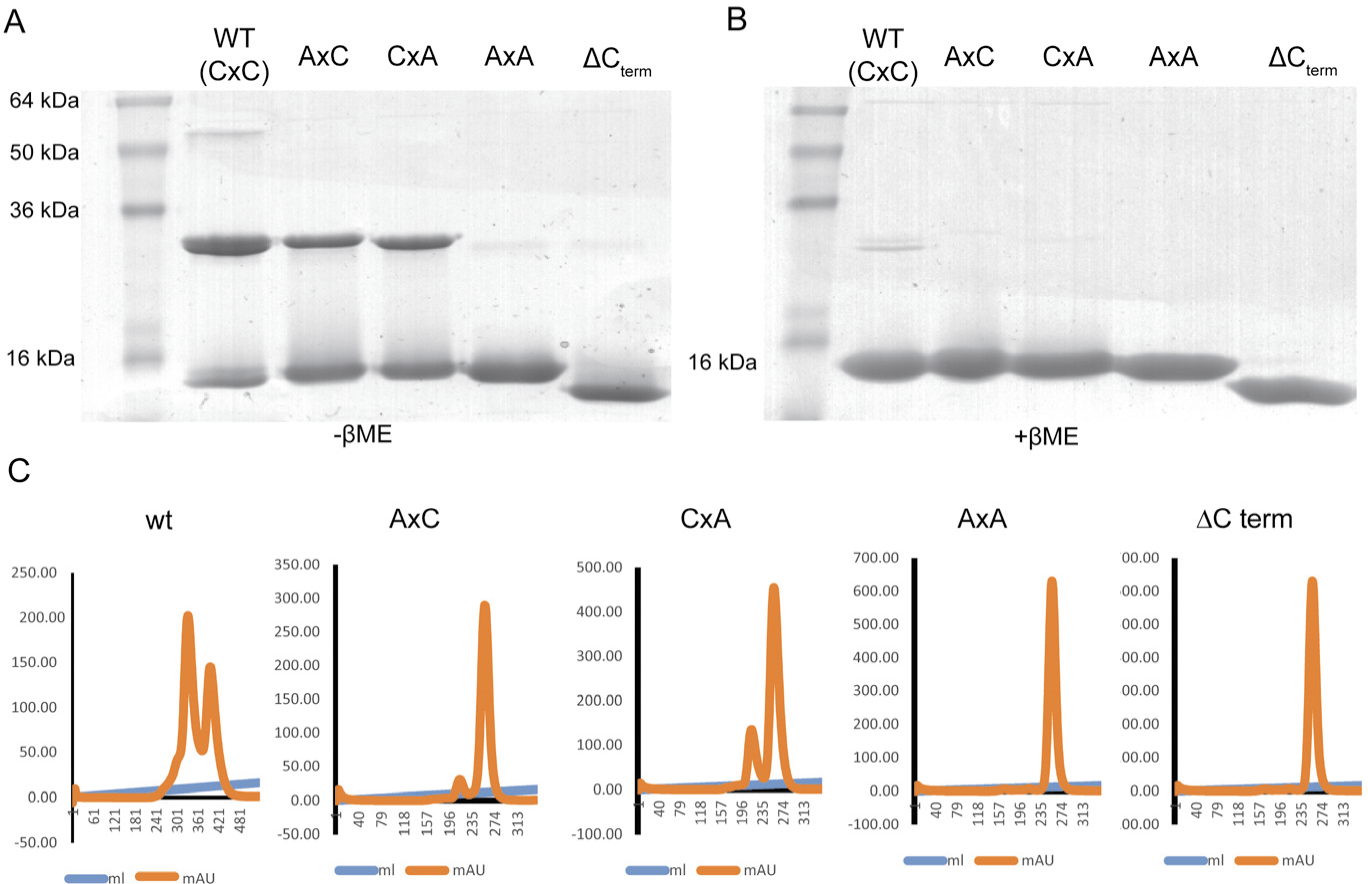
*In vitro* analysis of recombinant BslA higher order form. SDS-PAGE analysis of 30 μg of BslA, BslAAxC, BslACxA, BslAAxA, BslA ΔCterm recombinant protein in the absence **(A)** and presence **(B)** of reducing agent. **(C)** Size exclusion chromatography (SEC) analysis of recombinant protein using a Superdex 75 10/300GL column in an unreduced state.

**Figure S2:**
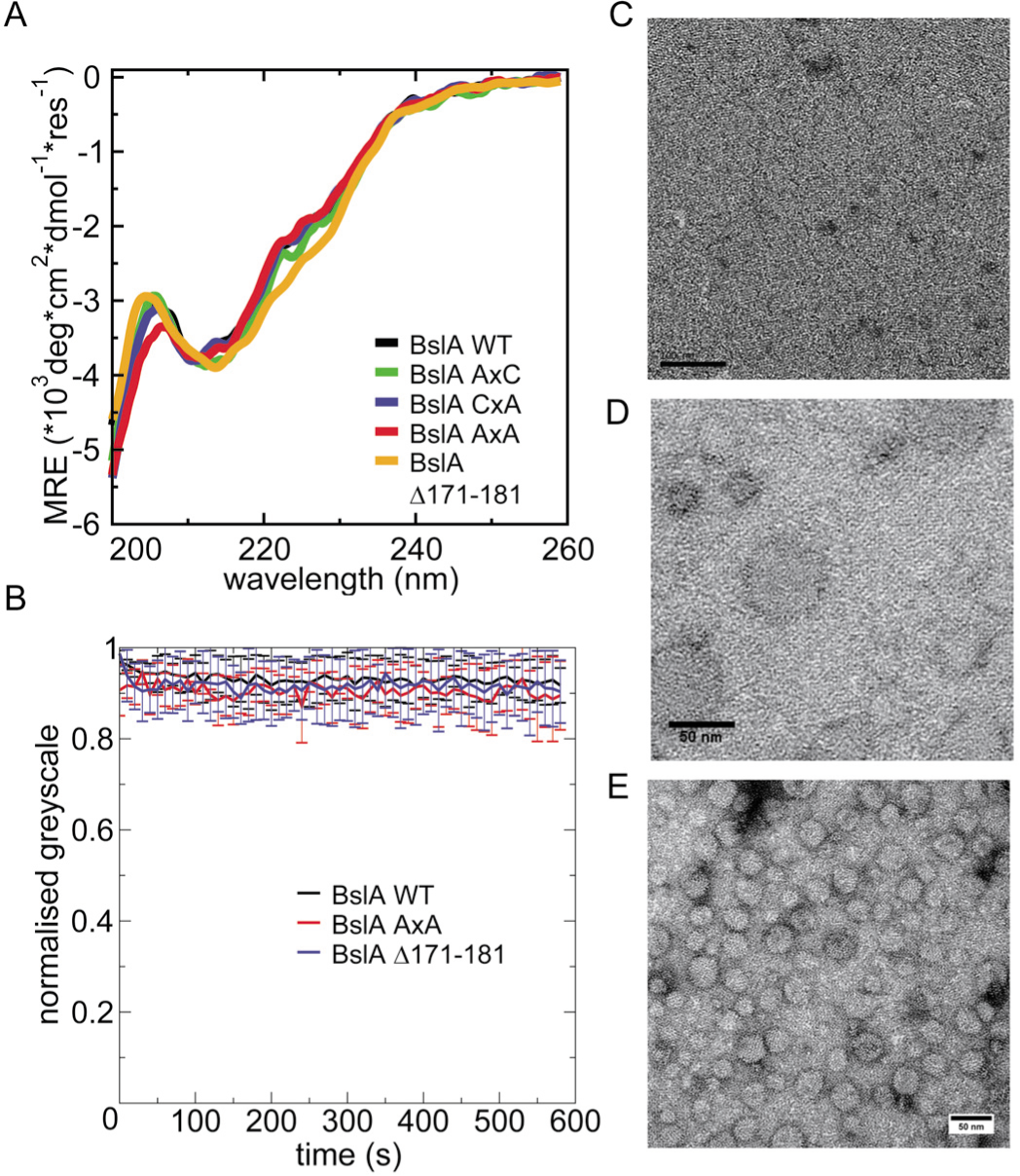
Presence of CxC motif does not affect *in vitro* characteristics of BslA. **(A)** Solution state circular dichroism spectra of recombinant BslA (black), BslA_C178A_ (green), BslA_C180A_(blue), BslA_C178A,C180A_ (red) and BslA_Δ171-181_ (orange) indicate all three variants have a similar secondary structure. **(B)** Wrinkle relaxation of WT BslA (black), BslA_C178A,C180A_ (red)and BslA_Δ171-181_ (blue). Wrinkles were formed in the protein film by removing fluid from the droplet and relaxation of the wrinkles was followed over time. The error bars represent the standard deviation in normalized greyscale over at least 5 independent wrinkles. Over ten minutes, wrinkles do not relax noticeably. Transmissionelectron microscopy images of BslA **(C)**, BslA_C178A,C180A_ **(D)** and BslA_Δ171-181_ **(E)** stained with uranyl acetate indicates that all three variants are capable of forming the ordered 2D lattice observed previously for WT BslA (11, 12).

**Figure S3:**
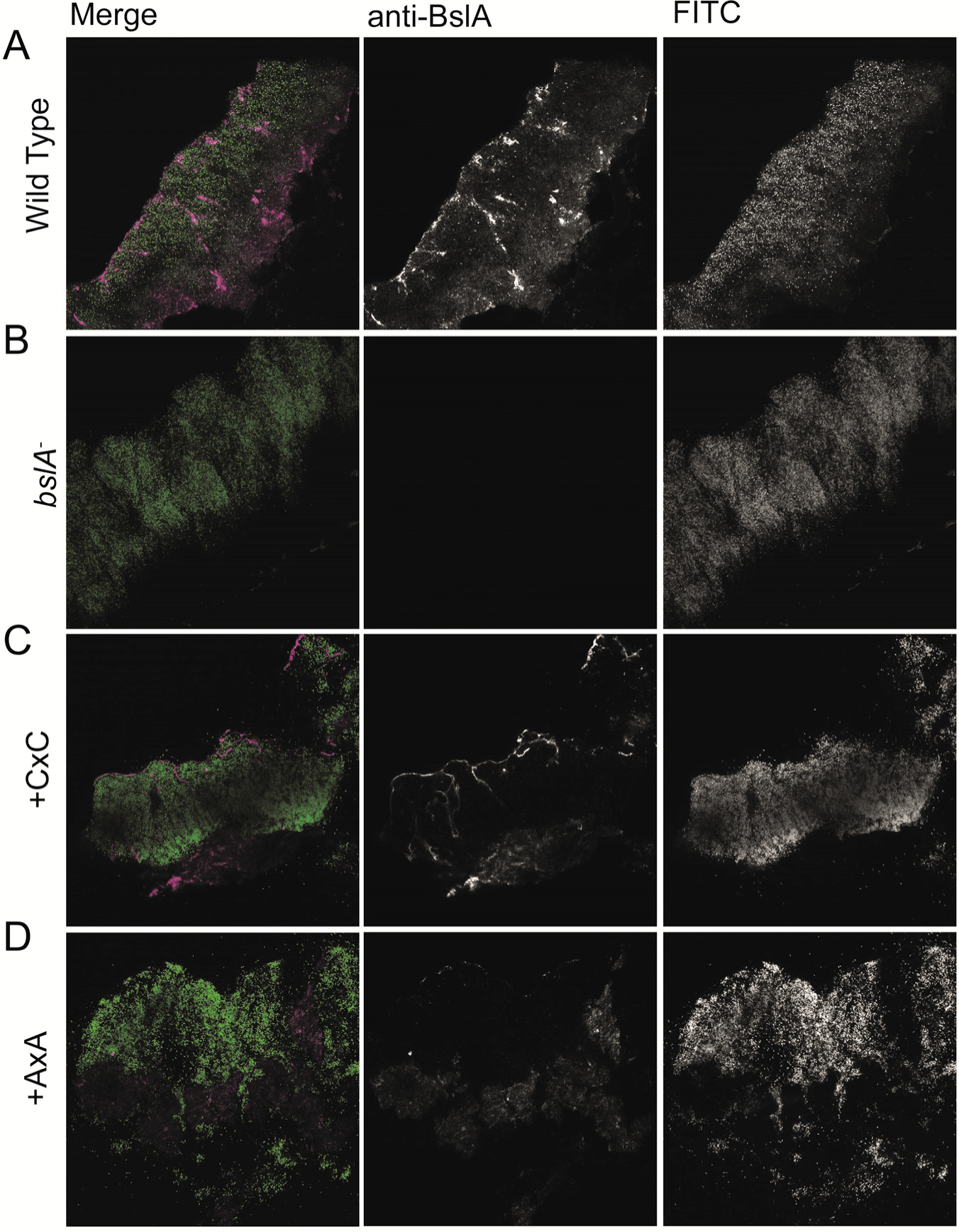
*In situ* analysis of BslA localisation in the biofilm. Confocal scanning laser microscopy images of cross sections through biofilms formed by **(A)** wild-type cells (NRS1473); **(B)** the *bslA* mutant strain (NRS3812); the *bslA* mutant complemented with either **(C)** the wild type variant of *bslA* (NRS5132); or**(D)** the gene encoding the BslA_AxA_ variant (NRS5136). For the merged images the fluorescence from the GFP within the cells is shown in green and the fluorescence associated with DyLight594, representing immuno-labelled BslA staining is shown in magenta. The scale bar represents 50 μm.

**Figure S4:**
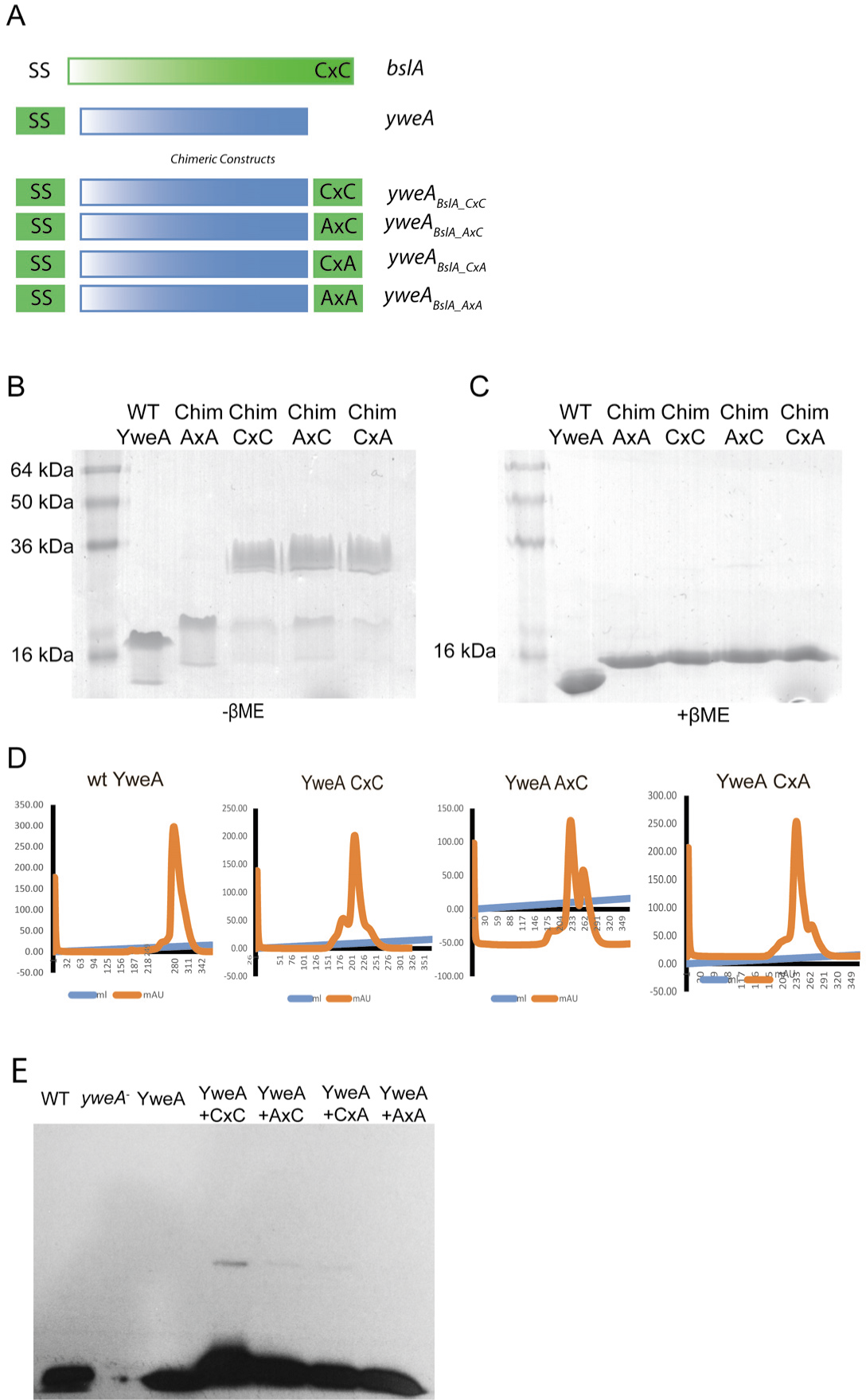
*In vitro* and *in vivo* analysis of recombinant YweA and chimeric YweA protein higher order forms. **(A)** Schematic of the YweA chimeric forms: ‘SS’ represents the BslA signal sequence used for export and ‘CxC’ the C-terminal 11 amino acids from BslA. SDS-PAGE analysis of 30 μg of YweA, YweA_BslA171-181_, YweA_BslA171-181 C178A_; YweA_BslA171-181 C180A_, YweA_BslA171-181 C178A C180A_ recombinant protein in the absence **(B)** and presence **(C)** of reducing agent. **(D)** Size exclusion chromatography (SEC) analysis of recombinant proteins using a Superdex75 10/300GL column in an unreduced state. **(E)** Western blot analysis of YweA in a reduced state detected from biofilm protein extracts.

**Figure S5:**
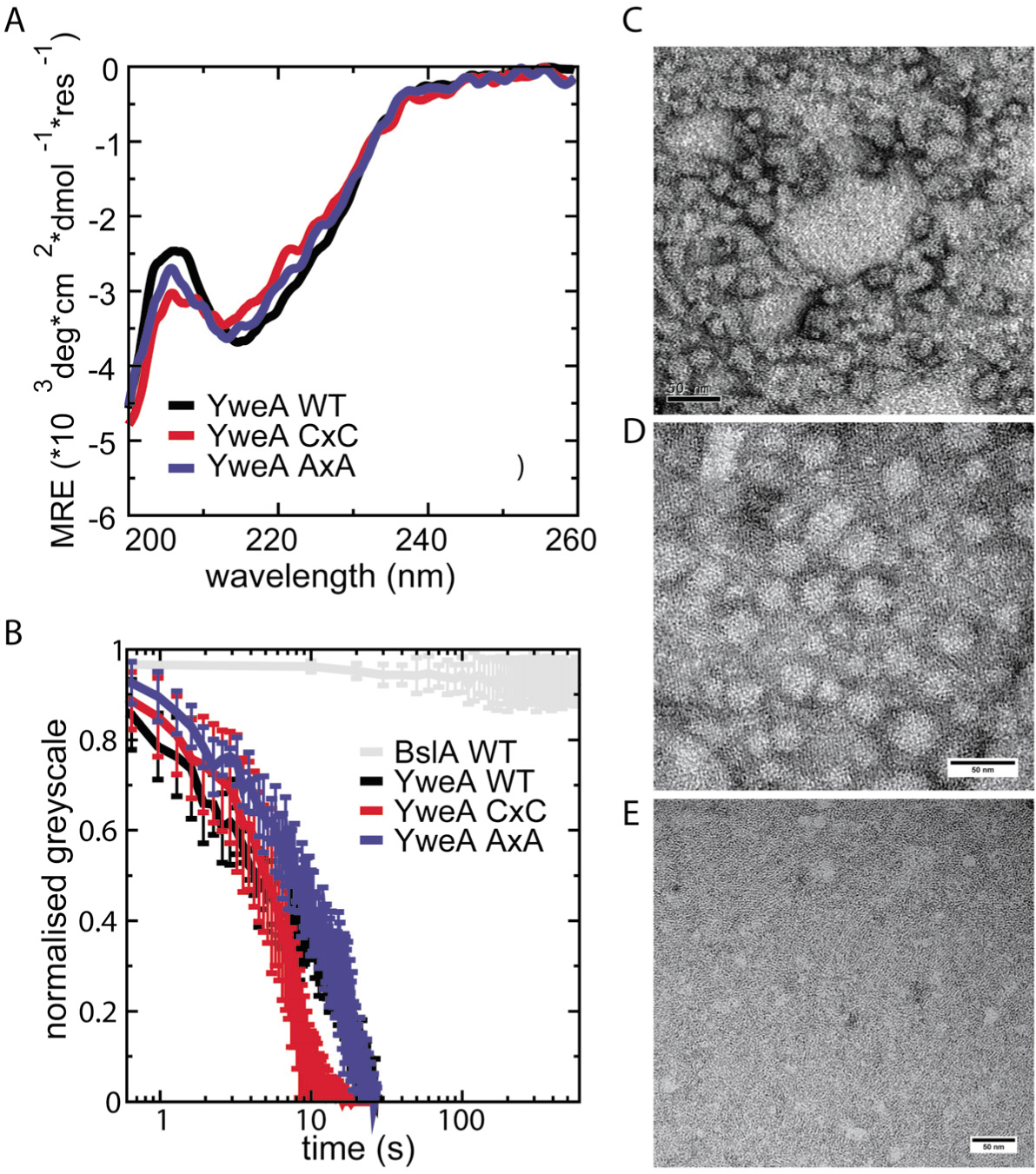
Transplantation of the CxC motif to the YweA C-terminus does not restore *in vitro* BslA behaviour. Solution state circular dichroism spectra of WT YweA (black), YweA_BslA171-181_ (red) andYweA_BslA171-181 C178A C180A_ (blue) indicate all three variants have a similar secondary structure. **(B)** Wrinkle relaxation of YweA (black), YweA_BslA171-181_ (red) and YweA_BslA171-181 C178A C180A_ (blue). Wrinkles were formed in the protein film by removing fluid from the droplet and relaxation of the wrinkles was followed over time. The error bars represent the standard deviation in normalized greyscale over at least 5 independent wrinkles. For all three variants, wrinkle relaxation is very rapid. Transmission electron microscopy images of YweA **(C)**, YweA_BslA171-181_ **(D)** and YweA_BslA171-181 C178A C180A_ **(E)** stained with uranyl acetate indicates that all three variants are capable of forming the ordered 2D lattice observed previously for WT BslA (11, 12) and YweA (44).

**Table S1:** Full list of strains used in this study.

| Strain | Relevant genotype /Description <sup>a</sup> | Source / Construction <sup>b,c</sup> |
| --- | --- | --- |
| *B. subtilis* |  |  |
| NCIB3610 | Prototroph | BGSC |
| 168 | *trpC2* | BGSC |
| NRS2097 | NCIB3610 *bslA::cml* | (45) |
| NRS2299 | NCIB3610 *bslA::cml amyE::Phy-spank-bslA-lacI (spc)* | (45) |
| NRS5165 | 168 *amyE::Phy-spank-bslA<sub>C178A</sub>-lacI (spc)* | pNW1500 → 168 |
| NRS5166 | 168 *amyE::Phy-spank-bslA<sub>C180A</sub>-lacI (spc)* | pNW1501 → 168 |
| NRS5167 | 168 *amyE::Phy-spank-bslA<sub>C178AC178A</sub>-lacI (spc)* | pNW1502 → 168 |
| NRS5177 | NCIB3610 *bslA::cml amyE::Phy-spank-bslA<sub>C178A</sub>-lacI (spc)* | SPP1 NRS5165 → NRS2097 |
| NRS5178 | NCIB3610 *bslA::cml amyE::Phy-spank-bslA<sub>C180A</sub>-lacI (spc)* | SPP1 NRS5166 → NRS2097 |
| NRS5179 | NCIB3610 *bslA::cml amyE::Phy-spank-bslA<sub>C178AC180A</sub>-lacI (spc)* | SPP1 NRS5167 → NRS2097 |
| NRS2953 | 168 *amyE::Phy-spank-bslA<sub>Δ172-181</sub>-lacI (spc)* | pNW621 → 168 |
| NRS2957 | NCIB3610 *bslA::cml amyE::Phy-spank-bslA<sub>Δ172-181</sub>-lacI (spc)* | SPP1 NRS2953 → NRS2097 |
| NRS1471 | 168 *sacA-Pspachy-gfp (kan)* | (12) |
| NRS1473 | NCIB3610 *sacA-Pspachy-gfp (kan)* | (12) |
| NRS3812 | NCIB3610 *bslA::cat, sacA::Pspachy-gfp (kan)* | (12) |
| NRS2404 | 168 *yweA::Kan* | (44) |
| NRS2405 | NCIB3610 *yweA::Kan* | (44) |
| NRS2409 | 168 *amyE::Phy-spank-yweA (spc)* | (44) |
| NRS2412 | NCIB3610 *bslA::cml amyE::Phy-spank-yweA (spc)* | (44) |
| NRS5550 | 168 *amyE::Phy-spank-bslAss-yweA (spc)* | pNW1719 → 168 |
| NRS5551 | NCIB3610 *bslA::cml amyE::Phy-spank-bslAss-yweA (spc)* | SPP1 NRS5550 → NRS2097 |
| NRS4831 | 168 *amyE::Phy-spank-bslAss-yweA. bslA<sub>171-181</sub> (spc)* | pNW1079 → 168 |
| NRS4834 | NCIB3610 *bslA::cml + amyE::Phy-spank-bslAss- yweA. bslA<sub>171-181</sub> (spc)* | SPP1 NRS4831 → NRS2097 |
| NRS5208 | 168 *amyE::Phy-spank-bslAss-yweA- bslA<sub>171-181 C178AC180A</sub>* | pNW1507 → 168 |
| NRS5210 | NCIB3610 *bslA::cml + amyE::Phy-spank-bslAss-yweA- bslA<sub>171-181 C178AC180A</sub>* | SPP1 NRS5208 → NRS2097 |
| NRS5509 | 168 *amyE::Phy-spank-bslAss-yweA- bslA<sub>171-181 C178A</sub>* | pNW1702 → 168 |
| NRS5515 | NCIB3610 *bslA::cml + amyE::Phy-spank-bslAss-yweA- bslA<sub>171-181 C178A</sub>* | SPP1 NRS5509 → NRS2097 |
| NRS5510 | 168 *amyE::Phy-spank-bslAss-yweA- bslA<sub>171-181 C180A</sub>* | pNW1703 → 168 |
| NRS5516 | NCIB3610 *bslA::cml + amyE::Phy-*<br>*spank-bslAss-yweA-bslA*<sub>171-181 C180A</sub> | SPP1 NRS5510→<br>NRS2097 |
| BKEE21460 | 168 *bdbA::erm* | BGSC |
| NRS5552 | NCIB3610 *bdbA::erm* | SPP1 NRS5598 →<br>NCIB3610 |
| *bdbCD* | 168 *bdbCD::spc* | (21) |
| NRS5554 | NCIB3610 *bdbCD::spc* | SPP1 NRS5139→<br>NCIB3610 |
| NRS5553 | NCIB3610 *bdbCD::spc bdbA::erm* | SPP1 NRS5139→<br>NRS5552 |
| NRS5132 | NCIB3610 *bslA::cml amyE-Pspac-hy-*<br>*gfpmut2 (spc) sacA::PbslA-bslA (kan)* | SSP1 NRS1113→NRS2978 |
| NRS5136 | NCIB3610 *bslA::cml amyE-Pspac-hy-*<br>*gfpmut2 (spc) sacA::PbslA-bslA*<sub>C178A C180A</sub><br>(*kan*) | SSP1 NRS5149→NRS5131 |
| *E. coli* |  |  |
| MC1061 | *E. coli F'lacIQ lacZM15 Tn10 (tet)* | *E. coli* Genetic Stock<br>Centre |
| BL21 (DE3) | F– *ompT hsdSB(rB–, mB–) gal dcm*<br>(DE3) | (46) |

- a. Drug resistance cassettes are indicated as follows: *cml*, chloramphenicol resistance; *kan*, kanamycin resistance; *mls*, lincomycin and erythromycin resistance and *spc*, spectinomycin resistance. BSGC represents the *Bacillus* genetic stock centre. - b. The direction of strain construction is indicated with DNA or phage (SPP1) (→) recipient strain. The reference is provided if the strain has previously been described.

**Table S2.**
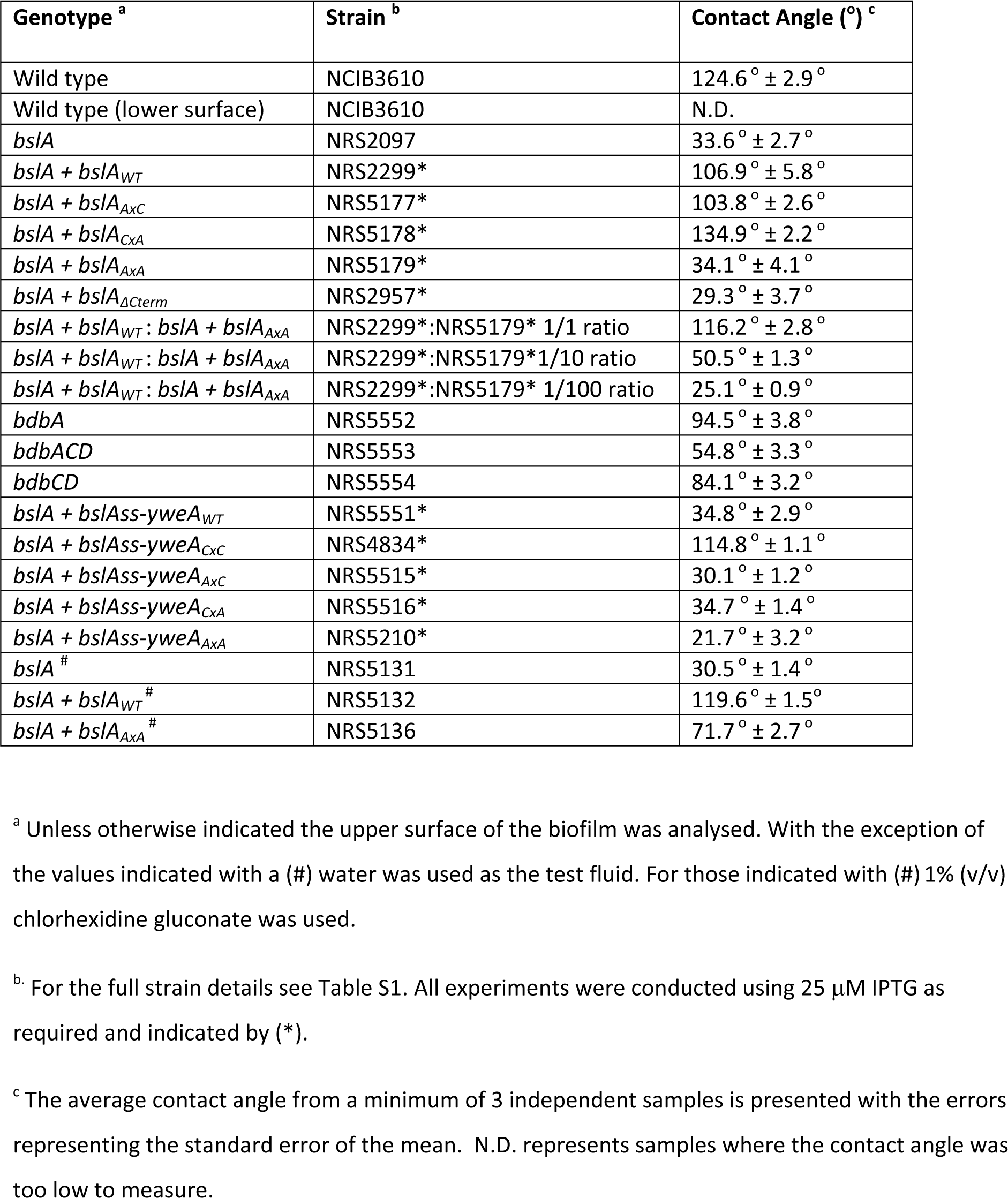
Contact angles of droplets placed on B. subtilis biofilms

**Table S3.**
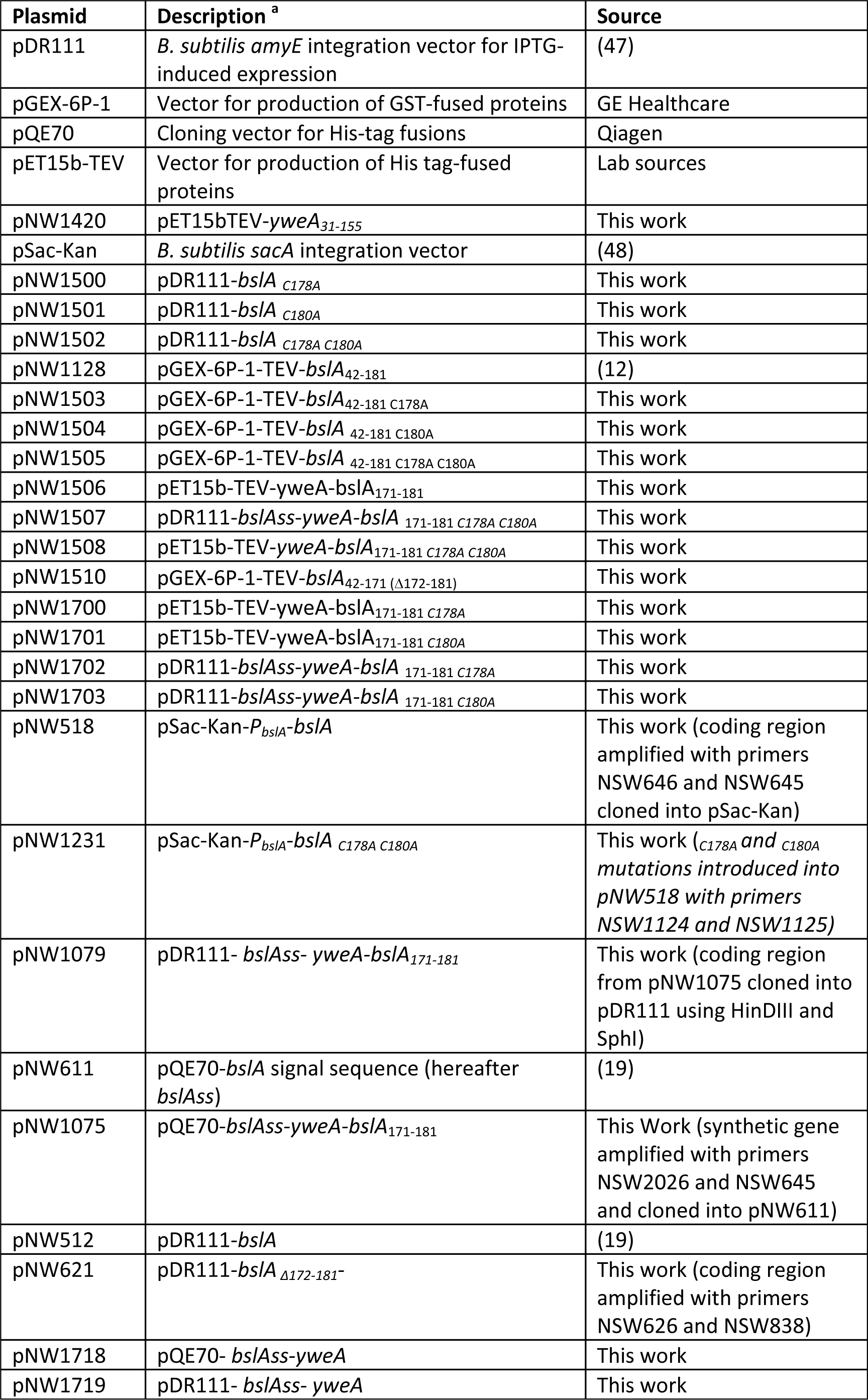
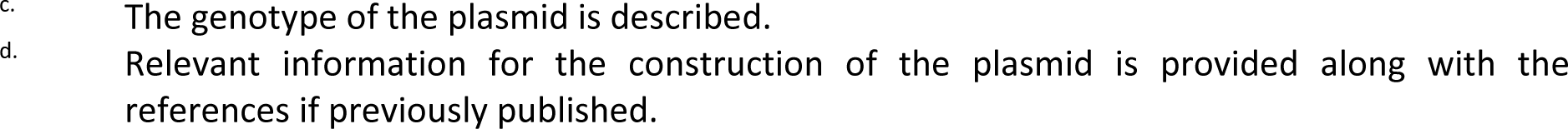
Plasmids used in this study

**Table S4:**
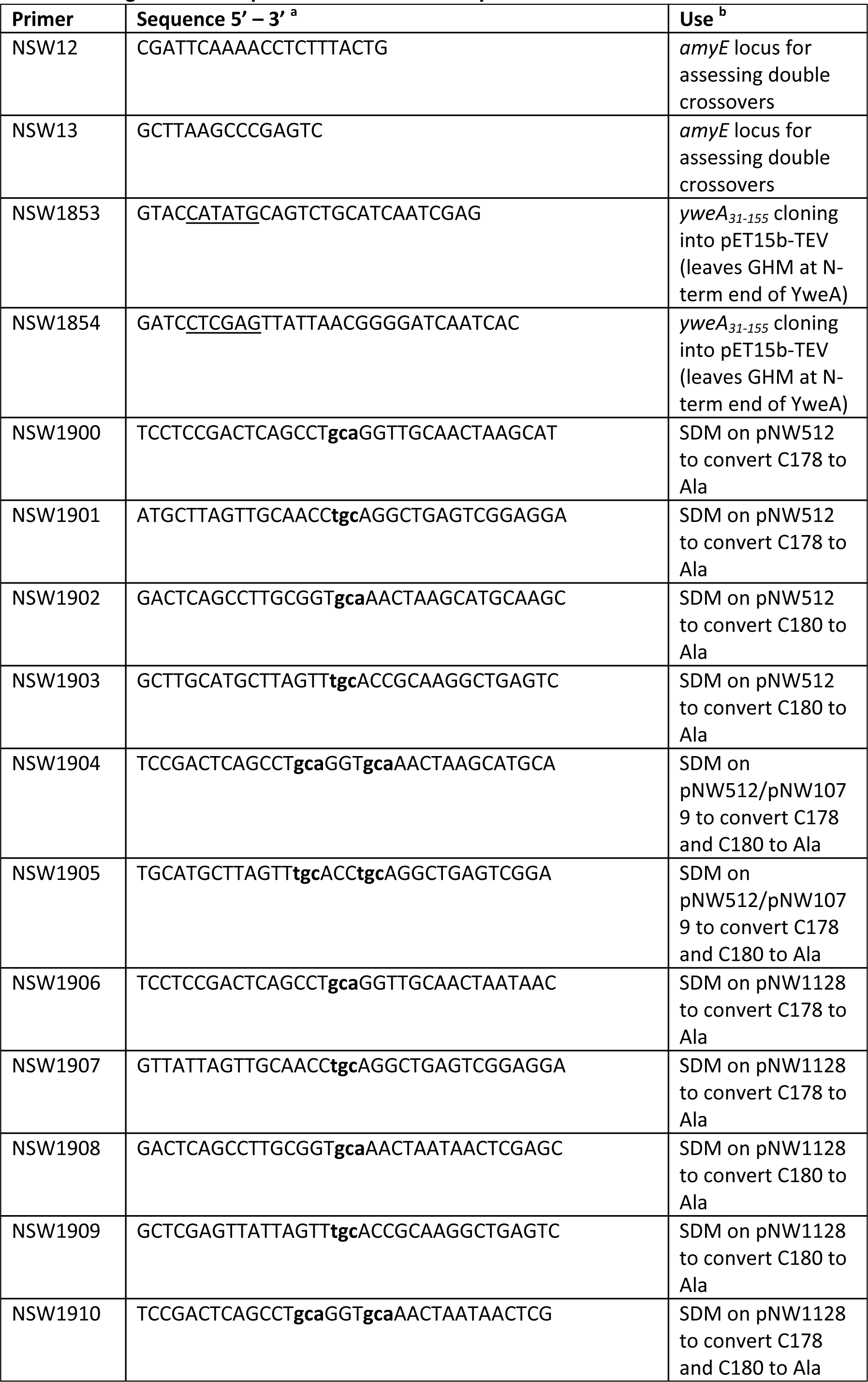
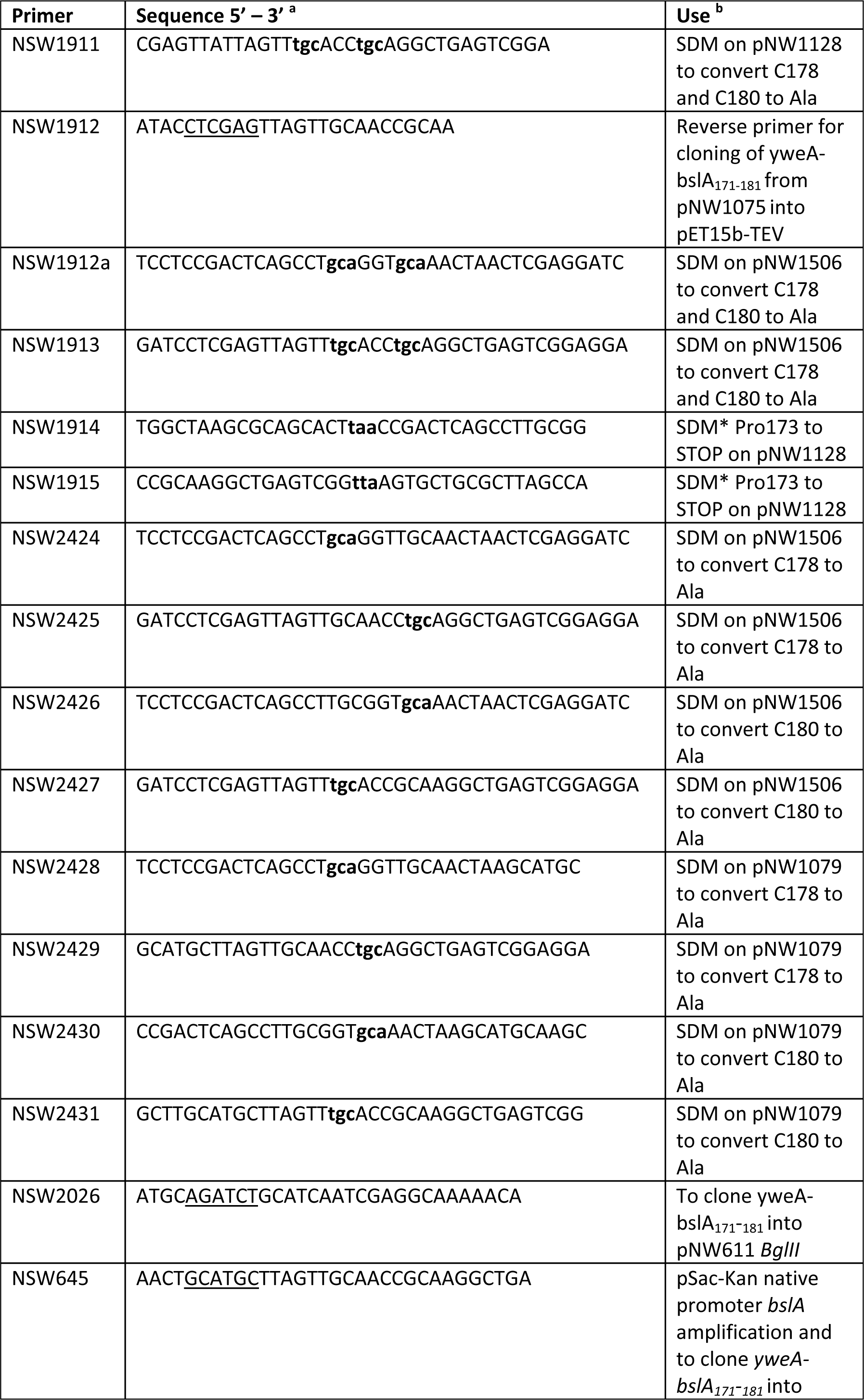
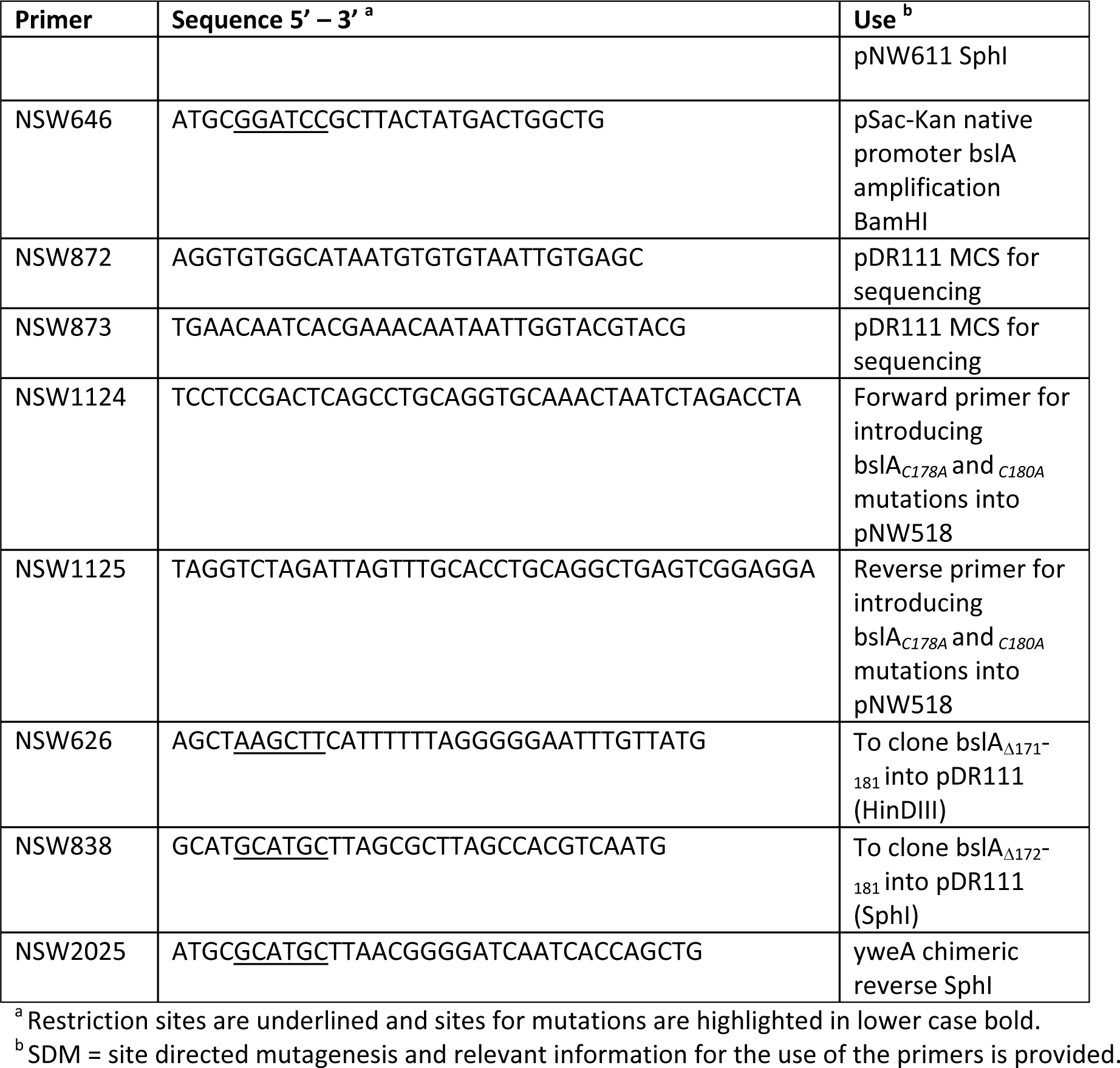
Oligonucleotide primers used in this study.

## References

1. Costerton JW, et al. (1987) Bacterial biofilms in nature and disease. Annu Rev Microbiol 41:435–464.

2. Flemming HC, et al. (2016) Biofilms: an emergent form of bacterial life. Nat Rev Microbiol 14(9):563–575.

3. Hobley L, Harkins C, MacPhee CE, & Stanley-Wall NR (2015) Giving structure to the biofilm matrix: an overview of individual strategies and emerging common themes. FEMS Microbiol Rev 39(5):649–669.

4. Earl AM, Losick R, & Kolter R (2008) Ecology and genomics of Bacillus subtilis. Trends Microbiol 16(6):269–275.

5. Branda SS, Gonzalez-Pastor JE, Ben-Yehuda S, Losick R, & Kolter R (2001) Fruiting body formation by Bacillus subtilis. Proc Natl Acad Sci U S A 98(20):11621–11626.

6. Epstein AK, Pokroy B, Seminara A, & Aizenberg J (2011) Bacterial biofilm shows persistent resistance to liquid wetting and gas penetration. P Natl Acad Sci USA 108(3):995–1000.

7. Arnaouteli S, MacPhee CE, & Stanley-Wall NR (2016) Just in case it rains: building a hydrophobic biofilm the Bacillus subtilis way. Curr Opin Microbiol 34:7-12.

8. Kobayashi K & Iwano M (2012) BslA (YuaB) forms a hydrophobic layer on the surface of Bacillus subtilis biofilms. Mol Microbiol 85(1):51–66.

9. Branda SS, Chu F, Kearns DB, Losick R, & Kolter R (2006) A major protein component of the Bacillus subtilis biofilm matrix. Mol Microbiol 59(4):1229–1238.

10. Romero D, Aguilar C, Losick R, & Kolter R (2010) Amyloid fibers provide structural integrity to Bacillus subtilis biofilms. Proc Natl Acad Sci U S A 107(5):2230–2234.

11. Bromley KM, et al. (2015) Interfacial self-assembly of a bacterial hydrophobin. Proc Natl Acad Sci U S A 112(17):5419–5424.

12. Hobley L, et al. (2013) BslA is a self-assembling bacterial hydrophobin that coats the Bacillus subtilis biofilm. P Natl Acad Sci USA 110(33):13600–13605.

13. Brandani GB, et al. (2015) The Bacterial Hydrophobin BslA is a Switchable Ellipsoidal Janus Nanocolloid. Langmuir 31(42):11558–11563.

14. Morris RJ, et al. (2016) Evolutionary variations in the biofilm-associated protein BslA from the genus Bacillus.

15. Poole LB (2015) The basics of thiols and cysteines in redox biology and chemistry. Free radical biology & medicine 80:148-157.

16. Kobashi K (1968) Catalytic oxidation of sulfhydryl groups by o-phenanthroline copper complex. Biochim Biophys Acta 158(2):239–245.

17. Bhushan B, Jung YC, & Koch K (2009) Micro-, nano-and hierarchical structures for superhydrophobicity, self-cleaning and low adhesion. Philosophical transactions. Series A, Mathematical, physical, and engineering sciences 367(1894):1631–1672.

18. Werb M, et al. (2017) Surface topology affects wetting behavior of Bacillus subtilis biofilms. Biofilms and Microbiomes 3(11).

19. Ostrowski A, Mehert A, Prescott A, Kiley TB, & Stanley-Wall NR (2011) YuaB functions synergistically with the exopolysaccharide and TasA amyloid fibers to allow biofilm formation by Bacillus subtilis. J Bacteriol 193(18):4821–4831.

20. Davey L, Halperin SA, & Lee SF (2016) Thiol-Disulfide Exchange in Gram-Positive Firmicutes. Trends Microbiol 24(11):902–915.

21. Kouwen TR, et al. (2007) Thiol-disulphide oxidoreductase modules in the low-GC Gram-positive bacteria. Mol Microbiol 64(4):984–999.

22. Crow A, et al. (2009) Crystal structure and biophysical properties of Bacillus subtilis BdbD. An oxidizing thiol:disulfide oxidoreductase containing a novel metal site. J Biol Chem 284(35):23719–23733.

23. Kobayashi K (2007) Gradual activation of the response regulator DegU controls serial expression of genes for flagellum formation and biofilm formation in Bacillus subtilis. Mol Microbiol 66(2):395–409.

24. Chai Y, Chu F, Kolter R, & Losick R (2008) Bistability and biofilm formation in Bacillus subtilis.Mol Microbiol 67(2):254–263.

25. Hogg PJ (2003) Disulfide bonds as switches for protein function. Trends Biochem Sci 28(4):210–214.

26. Schor M, Reid JL, MacPhee CE, & Stanley-Wall NR (2016) The Diverse Structures and Functions of Surfactant Proteins. Trends Biochem Sci 41(7):610–620.

27. Daniels R, et al. (2010) Disulfide bond formation and cysteine exclusion in gram-positive bacteria. J Biol Chem 285(5):3300–3309.

28. Meima R, et al. (2002) The bdbDC operon of Bacillus subtilis encodes thiol-disulfide oxidoreductases required for competence development. J Biol Chem 277(9):6994–7001.

29. Dorenbos R, et al. (2002) Thiol-disulfide oxidoreductases are essential for the production of the lantibiotic sublancin 168. J Biol Chem 277(19):16682–16688.

30. Oman TJ, Boettcher JM, Wang H, Okalibe XN, & van der Donk WA (2011) Sublancin is not a lantibiotic but an S-linked glycopeptide. Nat Chem Biol 7(2):78–80.

31. Nakamoto H & Bardwell JC (2004) Catalysis of disulfide bond formation and isomerization in the Escherichia coli periplasm. Biochim Biophys Acta 1694(1-3):111–119.

32. Santegoeds CM, Ferdelman TG, Muyzer G, & de Beer D (1998) Structural and functional dynamics of sulfate-reducing populations in bacterial biofilms. Appl Environ Microbiol 64(10):3731–3739.

33. Veach AM, Stegen JC, Brown SP, Dodds WK, & Jumpponen A (2016) Spatial and successional dynamics of microbial biofilm communities in a grassland stream ecosystem. Mol Eco l25(18):4674–4688.

34. Elias S & Banin E (2012) Multi-species biofilms: living with friendly neighbors. FEMS Microbiol Rev 36(5):990–1004.

35. Kempes CP, Okegbe C, Mears-Clarke Z, Follows MJ, & Dietrich LE (2014) Morphological optimization for access to dual oxidants in biofilms. Proc Natl Acad Sci U S A 111(1):208–213.

36. Okegbe C, Price-Whelan A, & Dietrich LE (2014) Redox-driven regulation of microbial community morphogenesis. Curr Opin Microbiol 18:39–45.

37. Smith DR, et al. (2015) In situ proteolysis of the Vibrio cholerae matrix protein RbmA promotes biofilm recruitment. Proc Natl Acad Sci U S A 112(33):10491–10496.

38. Serra DO, Richter AM, Klauck G, Mika F, & Hengge R (2013) Microanatomy at cellular resolution and spatial order of physiological differentiation in a bacterial biofilm. mBio 4(2):e00103–00113.

39. Harwood CR & Cutting SM (1990) Molecular biological methods for Bacillus. John Wiley & Sons Ltd. Chichester, England.

40. Verhamme DT, Kiley TB, & Stanley-Wall NR (2007) DegU co-ordinates multicellular behaviour exhibited by Bacillus subtilis. Mol Microbiol 65(2):554–568.

41. Hobley L, et al. (2013) BslA is a self-assembling bacterial hydrophobin that coats the Bacillus subtilis biofilm. Proceedings of the National Academy of Sciences of the United States of America 110(33):13600–13605.

42. Lee GF, Lebert MR, Lilly AA, & Hazelbauer GL (1995) Transmembrane signaling characterized in bacterial chemoreceptors by using sulfhydryl cross-linking in vivo. Proceedings of the National Academy of Sciences of the United States of America 92(8):3391–3395.

43. Lee GF, Burrows GG, Lebert MR, Dutton DP, & Hazelbauer GL (1994) Deducing the organization of a transmembrane domain by disulfide cross-linking. The bacterial chemoreceptor Trg. The Journal of biological chemistry 269(47):29920–29927.

44. Morris RJ, et al. (2016) The Conformation of Interfacially Adsorbed Ranaspumin-2 is an Arrested State on the Unfolding Pathway. arXiv:1602.04099.

45. J BacteriolMol MicrobiolVerhamme DT, Murray EJ, & Stanley-Wall NR (2009) DegU and Spo0A jointly control transcription of two loci required for complex colony development by Bacillus subtilis. J Bacteriol 191(1):100–108.

46. Studier FW & Moffatt BA (1986) Use of bacteriophage T7 RNA polymerase to direct selective high-level expression of cloned genes. Journal of molecular biology 189(1):113–130.

47. Britton RA, et al. (2002) Genome-wide analysis of the stationary-phase sigma factor (Sigma-H) regulon of Bacillus subtilis. J Bacteriol 184(17):4881–4890.

48. Middleton R & Hofmeister A (2004) New shuttle vectors for ectopic insertion of genes into Bacillus subtilis. Plasmid 51(3):238–245.

